# Host HSP90 mimicry by a parasite effector triggers mitochondrial membrane shedding

**DOI:** 10.64898/2026.11.16.766804

**Authors:** Julian Straub, Sebastian Kreimendahl, Jan-Hannes Schaefer, Sachin Yuvaraj Dinesh Babu, Fabian den Brave, Amy Prichard, Melissa Heredia, Kin W Chao, Katharina Stillger, Hina Ojha, Miratul M. K. Muqit, Alexey Amunts, Thomas Becker, Gabriel Lander, Sarah L. Rouse, Danielle A. Grotjahn, Lena Pernas

## Abstract

The outer mitochondrial membrane mediates critical host defenses against intracellular pathogens. How microbes manipulate this compartment remains poorly understood. Here, we found that the human parasite *Toxoplasma gondii* remodels host mitochondrial membranes through molecular mimicry of HSP90 chaperones. Cryo-electron microscopy analyses showed that the parasite effector TgMAF1 binds the same specialized clamp domain of the mitochondrial import receptor TOM70 as chaperones of the HSP70/90 family. Biochemical and comparative genomics analyses demonstrate that this mimicry arose through the neofunctionalization of the TgMAF1 C-terminus from an ancestral paralog incapable of TOM70 binding. In molecular dynamics simulations, complexed TgMAF1-TOM70 triggered the formation of higher order assemblies that induced local membrane curvature. These findings uncover a previously unknown host subversion strategy and establish molecular mimicry of a chaperone as a mechanism for pathogen-driven remodeling of the host OMM.

## Introduction

Mitochondria are dynamic organelles that integrate cellular and environmental cues to maintain homeostasis and coordinate cellular responses to stress ^1^. Their role in signal transduction extends to infection, during which mitochondria detect intracellular pathogens and activate innate immune defenses ^2–5^. Central to this function is the outer mitochondrial membrane (OMM), the interface between mitochondria and the cell that transduces cytosolic signals into changes in mitochondrial shape and function that mediate cellular responses ^1^. For example, the OMM protein MAVS translates the sensing of viral RNA into mitochondrial elongation and the activation of antiviral immunity ^6–8^. Consequently, viruses such as hepatitis C and SARS-CoV-2 have evolved effectors that cripple the MAVS-signaling axis by cleaving MAVS off the OMM or inhibiting its downstream adaptor TOM70, thereby preventing the induction of antiviral responses ^9, 10^. By contrast, how pathogens manipulate the architecture of the OMM remains little understood.

To address this question, we focused on the human parasite *Toxoplasma gondii* because of its ability to interact with and remodel the outer membrane of host mitochondria. After invasion, *Toxoplasma* tethers host mitochondria within 17 nm of its parasite vacuole (PV) to establish mitochondria-*Toxoplasma* PV contact sites termed host mitochondrial association (HMA) ^11, 12^. HMA requires the parasite effector protein TgMAF1 that is anchored to the PV membrane (PVM), and the host OMM receptor TOM70 ^13–15^. The single loss of either TgMAF1 or TOM70 prevents HMA ^13–15^.

Host mitochondria tethered at the *Toxoplasma* PV become highly enriched for TOM70 and release large micron-sized <u>s</u>tructures <u>p</u>ositive for <u>o</u>uter mitochondrial membrane termed SPOTs ^15^. Because the loss of TOM70 impairs parasite growth and OMM proteins MFN1 and MFN2 that mediate mitochondrial restriction of fatty acids from *Toxoplasma* are redistributed to SPOTs, SPOT induction is considered a parasite strategy to counter host defenses ^15–17^. Yet, how TgMAF1 hijacks TOM70 to remodel the OMM is unknown.

Here, we present the structure of human TOM70 in complex with TgMAF1. We found that TgMAF1 uses a C-terminal MLD motif to bind the clamp domain of TOM70, which docks the C-terminal EEVD motif of the HSP70/90 chaperone family that delivers preproteins to TOM70 for their import into mitochondria ^18–21^. A comparison of *TgMAF1* paralogs in *Toxoplasma* strains that do and do not tether mitochondria and subsequent biochemical analyses revealed that the TgMAF1 MLD motif is a functional mimic of the HSP70/90 EEVD motif. Investigating residues underlying the TgMAF1-TOM70 interaction revealed a previously unrecognized determinant of TOM70 binding to HSP70/90. In molecular dynamics simulations, complexed TgMAF1-TOM70 formed oligomeric assemblies that induced local membrane curvature. Thus, our findings indicate that TgMAF1 functions as a chaperone mimic and shed light on a mechanism by which an effector coopts TOM70 to remodel the host OMM.

### TgMAF1 traps TOM70 in an open state

The cytosolic domain of TOM70 consists of eleven tetratricopeptide repeats (TPR) that are arranged into three functionally distinct domains: the N-terminal clamp (TPR1-3), the core domain (TPR4-8), and the C-tail domain (TPR9-11) ^22–24^. The clamp domain docks HSP70/90 chaperones that deliver incoming mitochondrial preproteins, while the core and C-tail domains form a binding pocket that recognizes and engages preprotein substrates to enable their import ^18, 19, 24, 25^. Unbound TOM70 undergoes continuous sampling between a ‘closed’ conformation, in which the preprotein-binding cleft is occluded, and an open state that exhibits an accessible binding pocket (**Fig. 1a**) ^26^. This conformation sampling is thought to be allosterically coupled to chaperone engagement at the clamp, such that occupancy at one domain modulates the accessibility of the other ^26^.

**Figure 1:**
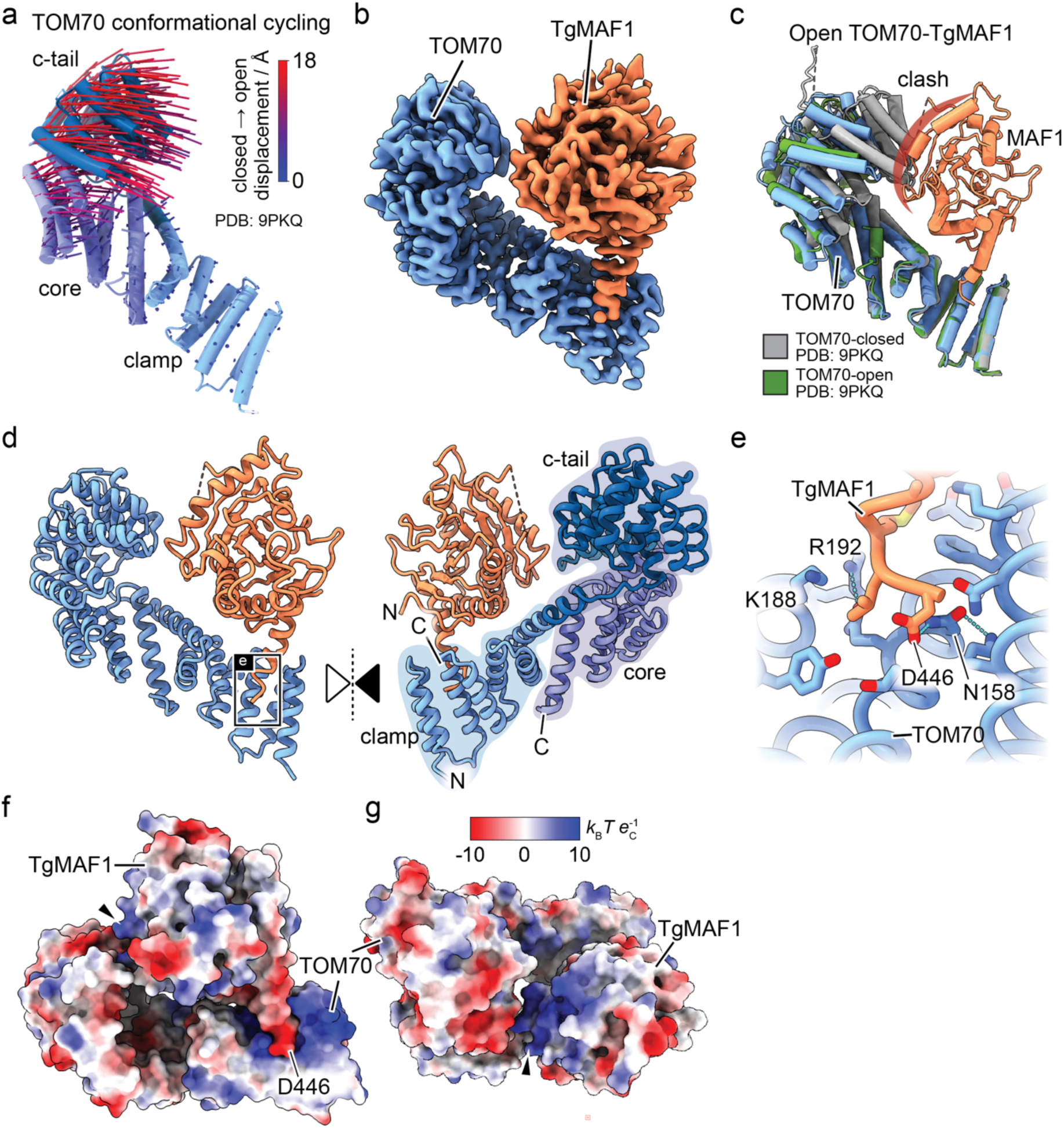
TgMAF1 traps TOM70 in an open state. **a,** Conformational cycling of TOM70 from closed to open using a vector displacement representation of PDB: 9PKQ. The TOM70 clamp, core and c-tail domains are annotated for orientation. **b,** Cryo-EM map of the TOM70-TgMAF1 complex. TOM70 and TgMAF1 are colored in blue and orange, respectively. **c,** Superimposition of the open TOM70-TgMAF1 (blue and orange, respectively) complex and closed TOM70 structure (PDB: 9PKQ, grey) with clash of MAF1 and TOM70 c-tail indicated in red. **d,** Atomic model of the complex with annotated termini. The inter-subunit contact is highlighted with a black box. **e**, Overview of subunit interactions with key residues mediating the contacts shown as sticks. Hydrogen bonds are shown as blue dotted lines. **f,g** Electrostatic surface potential from two viewing directions. TOM70-clamp interface to C-terminal TgMAF1, D446 of TgMAF1 annotated (e), and secondary binding site involving TOM70 c-tail, see Extended Data Fig. 6.

To elucidate the molecular basis of the interaction between TgMAF1 and TOM70, we determined the structure of the purified human TOM70-TgMAF1 complex using single-particle cryo-electron microscopy (cryo-EM). To do so, we co-expressed the cytosolic domain of human His_6_-tagged TOM70 (residues 111-608) and StrepII-tagged TgMAF1 (residues 173-446) in *E. coli* and assessed their interaction in bacterial lysates. TOM70^111–608^ was recovered in elutions of TgMAF1^WT^ as expected, given their interaction during infection (**Extended Data Fig. 1a-c**). The purified complex eluted as a single ∼50 kDa species, which showed monodisperse particles in negative-stain EM and resulting 2D projections, indicating stable complex formation (**Extended Data Fig. 1d-e, 2**). Initial single-particle analysis (SPA) revealed preferential orientation of the complex (**Extended Data Fig. 3, cryo-EM dataset 1**). Supplementing the sample with docyl maltoside (DM) during vitrification allowed us to resolve a consensus reconstruction of the complex at 3.4 Å global resolution (**Fig. 1b; Extended Data Fig. 3-4**) ^27^. While we could not completely remove anisotropy in our reconstruction, focused three-dimensional (3D) classification on MAF1 increased the local map quality, yielding a 3.4 Å reconstruction (**Extended Data Fig. 5**). Using AlphaFold3 (AF3) to guide our atomic modeling of the TgMAF1-TOM70 complex, we assigned densities for *Tg*MAF1^173-446^ and TOM70^111-604^; poorly resolved loops were built with low confidence, including TOM70^273-297^ and TgMAF1^275-281^, TgMAF1^367-372^ ^28^.

The overall architecture of the complex shows that TgMAF1 engages the cytosolic domain of TOM70, which adopted an extended conformation across its clamp, core, and C-tail domains (**Fig. 1b**). To determine which point along TOM70’s conformational continuum our complex represented, we superimposed our cryo-EM structure with the open and closed conformations of unliganded TOM70 ^26^. TgMAF1-bound TOM70 closely resembled the open TOM70 state; the open conformation accommodated the bound effector without overlap (**Fig. 1c**). However, modeling the closed conformation of TOM70 into our complex placed the C-tail in steric conflict with TgMAF1 (**Fig. 1c**). This indicates that TgMAF1 binding is incompatible with the closed state and selects for, or stabilizes, the open conformation of TOM70. Thus, in contrast to SARS-CoV-2 Orf9b that locks TOM70 in a partially closed, inhibited conformation, TgMAF1 traps TOM70 in an open state ^26^.

### TgMAF1 interacts with the chaperone-binding domain of TOM70

Our cryo-EM structure also revealed that TgMAF1 binds TOM70 through its C-terminal alpha-helix, which extends into the cradle-like pocket of the TOM70 clamp domain (**Fig. 1d**). This extended conformation positions the C-terminal Asp 446 of TgMAF1 deep within the TOM70 N-terminal clamp domain, explaining prior failures to detect an in vitro interaction with TOM70 when terminal TgMAF1 residues were truncated (**Fig. 1d-e; Extended Data Fig. 1a-c**) ^14^. This terminal residue is engaged through an electrostatically complementary interface: positively charged residues of TOM70, Arg 192 and Lys 188, form one side of the pocket, while on the opposite side TOM70 Asn 158 makes reciprocal hydrogen bonds with TgMAF1 Asp 446 (**Fig. 1e-h**). Adjacent to this pocket, the α4 helix of TgMAF1 engages α7 of TOM70 through polar and hydrophobic interactions mediated by TgMAF1 Leu 408, Thr 411, and TOM70 Gln 223 and Leu 226 (**Extended Data Fig. 6a-b**). We also observed a secondary, albeit less well-resolved, electrostatic interface between the negatively charged Asp 415 of TOM70 in TPR8 and Lys 268 of TgMAF1 (**Extended Data Fig. 6c**). Thus, TgMAF1 contacts TOM70 from head to tail, with its C-terminus buried in the clamp and a more N-terminal residue engaged at TPR8 of the TOM70 core domain.

To confirm the role of the TOM70 clamp domain in the interaction between TgMAF1 and TOM70 during infection, we leveraged the fact that TOM70 becomes highly enriched at the host mitochondria-*Toxoplasma* PV interface in a TgMAF1-dependent manner ^14, 15^. To this end, we compared the distribution of wild-type TOM70-GFP (TOM70^WT^) with that of TOM70 truncation mutants lacking the core and C-tail domain. We found that the TOM70 clamp domain (TOM70^1–235^) was sufficient to rescue the enrichment of TOM70 at ∼60% of *Toxoplasma* vacuoles, whereas the transmembrane domain (TMD) (TOM70^1–73^) alone was not (**Extended Data Fig. 6d-g**). Because our cryoEM analyses revealed a second interface between TgMAF1 and TPR 8 of the core domain of TOM70, we next expressed a TOM70 mutant lacking the C-tail TPRs (TOM70^1-477^). This mutant was sufficient to restore TOM70 enrichment to WT levels (**Fig. 1f-i**). In line with these results, a TOM70 mutant lacking the clamp domain (TOM70^Δclamp^) failed to enrich at mitochondria tethered to the PV (**Extended Data Fig. 7)**. Thus, the clamp domain of TOM70 is required for—and together with the core domain mediates—TOM70 sequestration at the host mitochondria-*Toxoplasma* PV interface.

### The C-terminal aspartate of TgMAF1 is critical for its binding to TOM70

To determine the TgMAF1 residues required for the enrichment of TOM70 at sites of HMA, we focused on the C-terminal TgMAF1 Asp 446 that inserts into the TOM70 clamp domain (**Fig. 1a-c**). We generated *Δmaf1 Toxoplasma* parasites expressing HA-tagged TgMAF1 carrying point mutants at Asp 446 and compared their ability to sequester TOM70 in infected cells. Unlike *Δmaf1:HA-MAF1^WT^*parasites, we found that *Δmaf1:HA-MAF1^D446Δ^* parasites were completely deficient for the perivacuolar enrichment of TOM70 (**Fig. 2a-b**). Similar results were observed during infection with *Δmaf1:MAF1^D446N^* and *Δmaf1:HA-MAF1^D446E^*parasites harboring either a structurally similar asparagine or negatively charged glutamate in lieu of aspartate (**Fig. 2a-b**). We next asked how the loss of TOM70 enrichment impacts host mitochondria-*Toxoplasma* PV contact sites. To do so, we performed electron microscopy analysis of HeLa cells infected with *Δmaf1* parasites expressing TgMAF1 point mutants. Whereas *Δmaf1:HA-MAF1^WT^*parasite vacuoles showed the expected formation of host mitochondria-PV contact sites, *Δmaf1: HA-MAF1^D446Δ^, Δmaf1: HA-MAF1^D446N^,* and *Δmaf1: HA-MAF1^D446E^* all lacked detectable contact **(Fig. 2c-d).** Thus, TgMAF1 Asp 446 is required for the enrichment of TOM70 at mitochondria tethered to the *Toxoplasma* PV and for the establishment of HMA.

**Figure 2:**
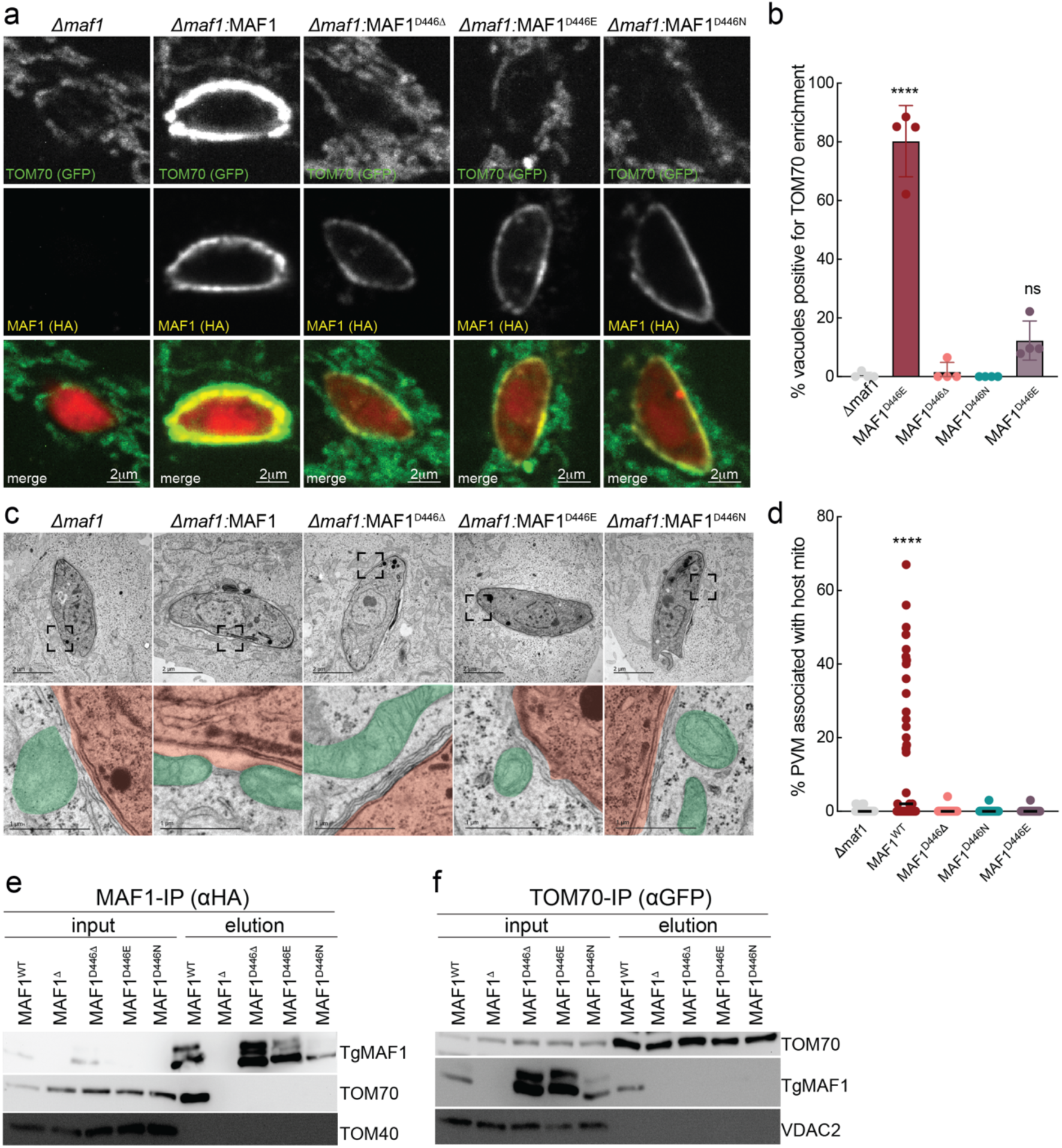
The C-terminal aspartate of TgMAF1 is required for TOM70 enrichment and the establishment of mitochondria-*Toxoplasma* contact sites. **a**, Immunofluorescence images of TOM70-GFP-expressing murine embryonic fibroblasts (MEFs) 5 hours post infection (hpi) with *Toxo-*mCh; scale bar 2 μm. (TgMAF1, HA; TOM70; GFP). **b,** Percentage of vacuoles positive for TOM70 enrichment in experiments as in a; >40 vacuoles for n=3 biological replicates; ****p<0.001 by one-way ANOVA, comparison to *Δmaf1*. **c,** Representative electron micrograph (EM) images of HeLa cells 5 hpi with *Toxoplasma* strains as in (a). **d,** Quantification of the percentage of the PVM in contact with host mitochondria from images as in (c), >20 vacuoles counted from n=1 biological replicate. **e,** Anti-HA IPs from WT Hela cells 24 hpi with *Δmaf1 Toxoplasma* parasites expressing the indicated HA-TgMAF1 mutants were analyzed by means of immunoblotting (IB) for TgMAF1 ∼60kDa, TOM70 ∼72 kDa; TOM40 ∼40kDa. (**f**) anti-GFP IPs from MEFs stably expressing TOM70-GFP 24 hpi with *Toxoplasma* strains as in (e) and analyzed by IB; VDAC2 ∼34 kDa.

Our results implicated Asp 446 as essential for TgMAF1 function but did not distinguish whether it mediated direct binding to TOM70 or acted indirectly. To test whether TgMAF1 binds TOM70 in an Asp 446-dependent manner, we immunopurified TgMAF1 from cells infected with *Δmaf1:HA-MAF1^D446Δ^, Δmaf1:HA-MAF1^D446N^, Δmaf1:HA-MAF1^D446E^* parasites. Immunoblot analysis revealed that TOM70 was enriched in immunoprecipitates of *HA-MAF1^WT^,* but not any of the point mutants (**Fig. 2e**). The reciprocal was also the case in infected cells: stably expressed TOM70-GFP co-immunoprecipitated TgMAF1^WT^, but not TgMAF1^D446Δ^, TgMAF1^D446E^, or TgMAF1^D446N^ (**Fig. 2f**). Thus, TgMAF1 Asp 446 is specifically recognized by the TOM70 clamp domain and is required for TOM70 binding.

### TgMAF1 is an HSP70/90 chaperone mimic

The interaction between the C-terminus of TgMAF1 and the clamp domain of TOM70 was reminiscent of the mechanism by which members of the HSP70/90 chaperone family interact with TPR proteins such as TOM70 ^18–21^. The C-terminal EEVD motif of HSP70/90 chaperones enables a two-carboxylate anchor with the positively charged TOM70 clamp, paralleling the interaction between the TgMAF1 C-terminus and the TOM70 clamp ^19^. To investigate the possibility that TgMAF1 functionally mimics HSP70/90 to bind TOM70, we superimposed our TgMAF1-TOM70 map with structures of HSC70 and HSP90 peptides bound to TPR1A and TPR2A of co-chaperone HOP, respectively. Analogous to the C-terminal region of HSP70/90 EEVD, TgMAF1 D446 forms hydrogen bonds with conserved basic residues within the clamp domain of TOM70, K188 and R192 (**Fig. 3a-b**)^18^. These data suggest that TgMAF1 evolved to mimic the canonical HSP70/90 EEVD-clamp binding mechanism.

**Figure 3:**
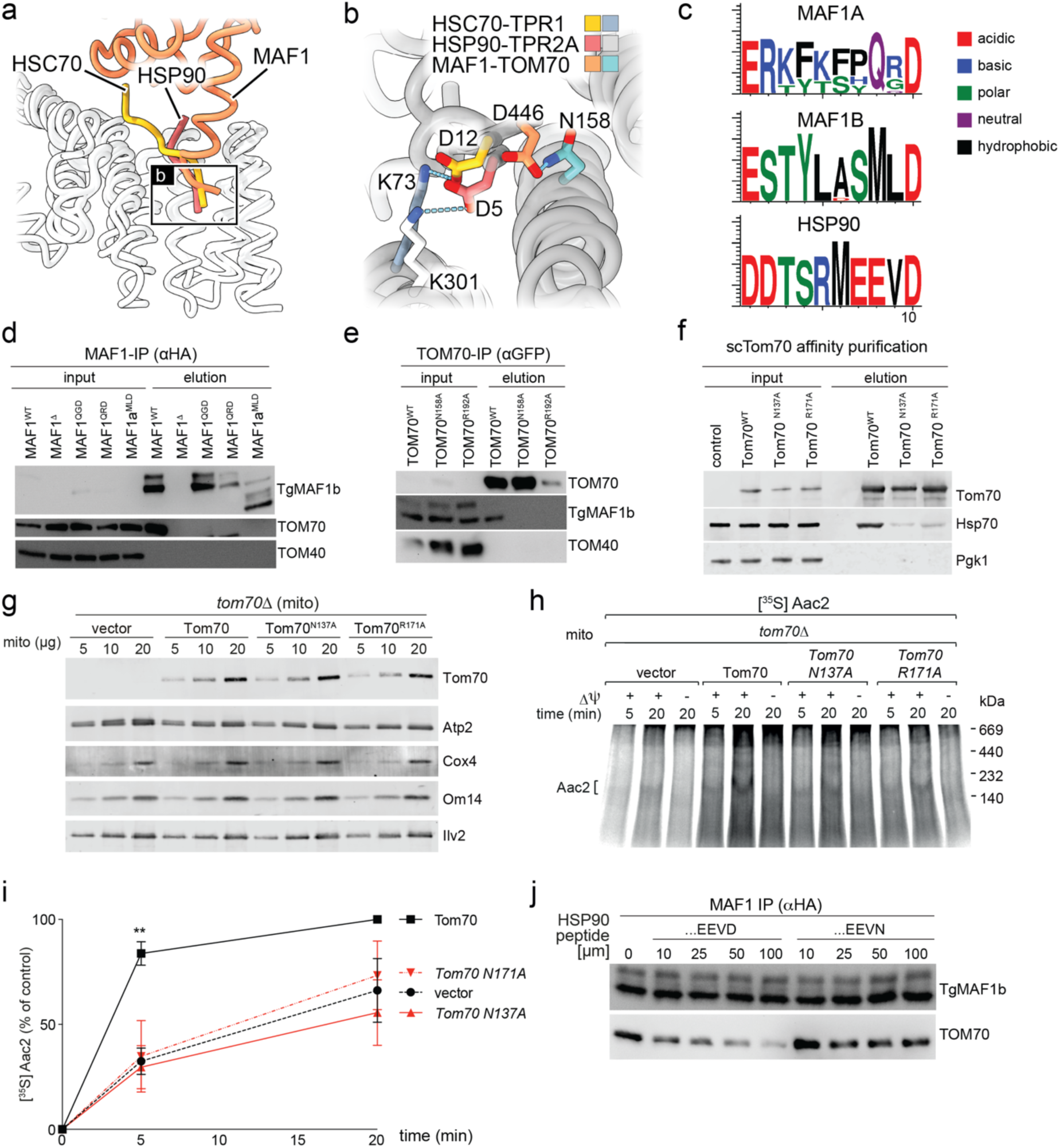
TgMAF1 mimics HSP90 to bind TOM70. **a,** Superimposed structures of the carboxyterminal region of MAF1-TOM70, HSC70-TPR1 (PDB: 1ELW) and HSP90-TPR2A (PDB: 1ELR). **b,** Binding mode of the carboxy-terminal Asp shown as sticks with H-bonds in cyan. **c,** Sequence logos (Weblogos) for TgMAF1a and TgMAF1b paralogs identified in the major *Toxoplasma* lineages and HSP90 from diverse model organisms (sequences in Ext. Data Fig. 9). The height of the amino acid stack is proportional to the sequence conservation at that position. **d,** Anti-HA IPs from WT Hela cells 24 hours post infection (hpi) with *Δmaf1* parasites expressing the indicated HA-tagged TgMAF1 mutants were analyzed by means of immunoblotting (IB); TOM70, ∼72kDa, TgMAF1(HA), ∼60kDa, TOM40, ∼40kDa. **e**, Anti-GFP IPs from HeLa cells stably expressing GFP fused to the indicated TOM70 variant 5 hpi with WT *Toxoplasma* were analyzed by IB; TgMAF1 ∼60 kDa; TOM70-GFP ∼105kdA, TOM40 ∼40kDa; VDAC2 ∼34 kDa. **f,** Yeast cytosolic extract was incubated with his-tagged TOM70 (cytosolic domain) and subjected to affinity purification with Ni-NTA agarose. Input and elution were analyzed by means of IB; Hsp70 ∼70 kDa; Tom70 ∼70 kDa; Pgk1 ∼55 kDA. **g,** Indicated amount (μg) of mitochondria isolated from indicated yeast stains were analyzed for indicated proteins by IB; Atp2 ∼55 kDa; Cox4 ∼15 kDa; Om14 ∼15kDa; Ilv2 ∼70 kDa. **h,** [^35^S]AAC import into mitochondria isolated from cells in (g) were analyzed by means of blue native gel and autoradiography. In control reactions, mitochondria were treated with valinomycin to deplete membrane potential (Δψ). **i,** Quantification of Aac2 import as in (h). Data are mean of the amount of imported AAC from n=5 independent experiments ±SEM, **p<0.01 for vector versus TOM70 by means of two-way ANOVA analysis. The amount of AAC for the longest import period in WT mitochondria was set to 100% **j,** Anti-HA IPs from WT Hela cells 24 hpi with *Δmaf1* parasites expressing HA-tagged TgMAF1 were performed in indicated concentrations of decapeptide corresponding to the last 10 residues of HSP90 and analyzed by means of IB.

To address this possibility, we leveraged the functional differences between TgMAF1, also referred to as TgMAF1b, and its paralog TgMAF1a that does not tether mitochondria ^14, 29^. Multiple tandemly duplicated copies of *TgMAF1a* and *TgMAF1b* that vary in sequence and copy number across *Toxoplasma* strains are encoded by the *TgMAF1* locus ^30^. Phylogenetic analyses indicate that *TgMAF1a* is the ancestral paralog, and that subsequent duplication and diversification gave rise to *TgMAF1b* as a mitochondrial tether ^29^. Indeed, unlike TgMAF1b, TgMAF1a has no predicted interactions with TOM70 according to AlphaFold (AF2) (**Extended Data Fig. 8**) ^14^.

We therefore asked whether the neofunctionalization of TgMAF1b as host mitochondrial tether coincided with the acquisition of residues resembling those found in HSP90. To address this, we compared the 10 C-terminal residues of TgMAF1a and TgMAF1b paralogs from diverse *Toxoplasma* lineages (**Extended Data Fig. 9**). Sequence logo analysis revealed that the positively charged lysines and arginines present in the TgMAF1a C-terminus were lost in TgMAF1b (**Fig. 3c; Extended Data Fig. 9**). This loss suggested a selection against basic residues in TgMAF1b and was consistent with our cryo-EM data showing electrostatic interactions between the negatively charged TgMAF1 (TgMAF1b) C-terminus and the positively charged TOM70 clamp domain (**Fig. 1d-e**).

We also observed that a MLD motif at the TgMAF1 C-terminus, which resembled the canonical EEVD found at the C-terminus of HSP90, was conserved across all homologs of TgMAF1b but not TgMAF1a (**Fig. 3c; Extended Data Fig. 9**)^19^. To test the importance of the MLD motif in TOM70 binding, we infected cells with *Δmaf1* parasites expressing TgMAF1 mutants harboring the corresponding *TgMAF1a* residues—glutamine and arginine (TgMAF1^QRD^) or glutamine and glycine (TgMAF1^QGD^) in lieu of Met 444 and Leu 445. Neither mutant co-immunoprecipitated with TOM70, nor led to the TOM70 enrichment at the host mitochondria-*Toxoplasma* PV interface (**Fig. 3d; Extended Data Fig. 10**). Of note, the expression of a TgMAF1a variant engineered to carry the C-terminal MLD motif also failed to interact with TOM70 (**Fig. 3d**). Thus, the MLD motif is essential, but not sufficient for TgMAF1b to bind and sequester TOM70.

If TgMAF1b engages TOM70 through a mechanism analogous to that of HSP90, we reasoned that residues critical for HSP90 binding would also be required for TgMAF1 binding and vice versa. To test this, we assessed TgMAF1 binding to the TOM70^R192A^ point mutant that is deficient for HSP70/90 binding, and thus in precursor protein import ^18^. As our cryo-EM structure revealed that TgMAF1 Asp 446 interacted with TOM70 158 Asn, we also tested a TOM70^N158A^ point mutant (**Fig. 1**). Following infection, TgMAF1 was recovered in elutions of TOM70 WT, but not TOM70^R192A^ nor TOM70^N158A^ (**Fig. 3e**). This result raised the question of whether TOM70 Asn 158 was critical for HSP70/90 binding.

TOM70 Asn158 was of particular interest because it is conserved in TPR domains of diverse proteins that interact with chaperones, including that of human (HsTOM70) and yeast (ScTom70), but whose contribution to chaperone binding had not yet been experimentally addressed ^18, 19^. To address this gap, we turned to baker’s yeast, a system in which ScTom70-Hsp70/90 chaperone interactions are well characterized. We found that Hsp70 bound to affinity-purified ScTom70, but not ScTom70 mutants harboring the R171A (HsTOM70 R192A) or N137A (HsTOM70 N158A) mutations (**Fig. 3f**). Furthermore, mitochondria isolated from these mutant yeast strains exhibited impaired import of the scTom70-dependent precursor Aac2 despite similar mitochondrial protein abundance, confirming a defect in chaperone delivery (**Fig. 3g-i**). Thus, analyzing how TgMAF1 engages TOM70 revealed that Asn 158, a previously uncharacterized residue, is critical for Hsp70/90 recognition.

Our results thus far supported a model in which *Toxoplasma* TgMAF1 hijacks TOM70 by mimicking HSP90. A logical prediction therefore was that the parasite effector and host HSP90 compete for TOM70 occupancy. To test this, we immunoprecipitated TgMAF1 from infected cells in the presence of increasing concentrations of a C-terminal decapeptide of HSP90, or a mutant version in which the EEVD motif was replaced with EEVN. We found that only the EEVD peptide diminished TgMAF1 binding to TOM70 at 10μM, and almost completely blocked binding at higher concentrations (**Fig. 3j**). Thus, TgMAF1 is a chaperone mimic that competes with HSP70/90 to bind TOM70. In line with these results, a structural superposition of our TOM70-TgMAF1 complex onto a high-confidence AF3-predicted TOM70-HSP90A complex confirmed the interaction between the HSP90A C-terminus and the TOM70 clamp domain, and revealed that helix α2 of HSP90A contacts TOM70 α7 in the same manner as TgMAF1 (**Extended Data Fig. 6a-c; Extended Data Fig. 11**).

A functional chaperone mimic should dock at the clamp domain while leaving the TOM70 preprotein-binding pocket accessible. To test whether TgMAF1-bound TOM70 could accommodate a preprotein, we turned to Orf9b, the SARS-CoV-2 effector that binds this pocket ^31, 32^. Rigid superposition of the TOM70-Orf9b and TgMAF1-TOM70 structures produced a steric clash between the TOM70 c-tail and TgMAF1 (**Extended Data Fig. 12a**). However, AF3 modeling of the TOM70-TgMAF1-Orf9b ternary complex predicted unimpaired Orf9b binding (**Extended Data Fig. 12a-c**). Consistent with the possibility that TgMAF1 and a preprotein can occupy TOM70 simultaneously, Orf9b co-enriched with TOM70 at TgMAF1-tethered mitochondria in *Toxoplasma-*infected cells expressing Orf9b (**Extended Data Fig. 12d**).

### Oligomerization of TgMAF1-TOM70 complexes and membrane curvature in molecular dynamics simulations

Having defined the structural mechanism driving the interaction between TgMAF1 and TOM70, we next asked how this interaction contributed to the remodeling of the OMM during infection. Crowding among membrane-bound proteins increases membrane curvature and drives membrane fission ^33^. TgMAF1 drives the marked accumulation of TOM70 on mitochondria tethered to the PV, the subpopulation of mitochondria from which SPOTs are released (**Fig. 1f-h**; **Fig. 2a**). We thus next addressed whether TgMAF1-TOM70 complexes aggregate in a manner that induces membrane bending. To do so, we performed 10 μs coarse-grained molecular dynamics simulations of full-length TgMAF1-TOM70 complexes embedded in representative PVM and OMM phospholipid bilayers in a 3×3 array, such that each bilayer contained nine transmembrane helices (TMHs) at the start (**Extended Data Fig. 13, Table 2-3**). By 10 μs, the nine complexed TgMAF1 and TOM70 TMHs coalesced into 2-3 assembled clusters while simulations with TgMAF1 or TOM70 alone showed 7-9 clusters (**Fig. 4a-b; Extended Data Fig. 13**). Thus, complexed TgMAF1-TOM70 nucleates higher order complex assembly.

**Figure 4:**
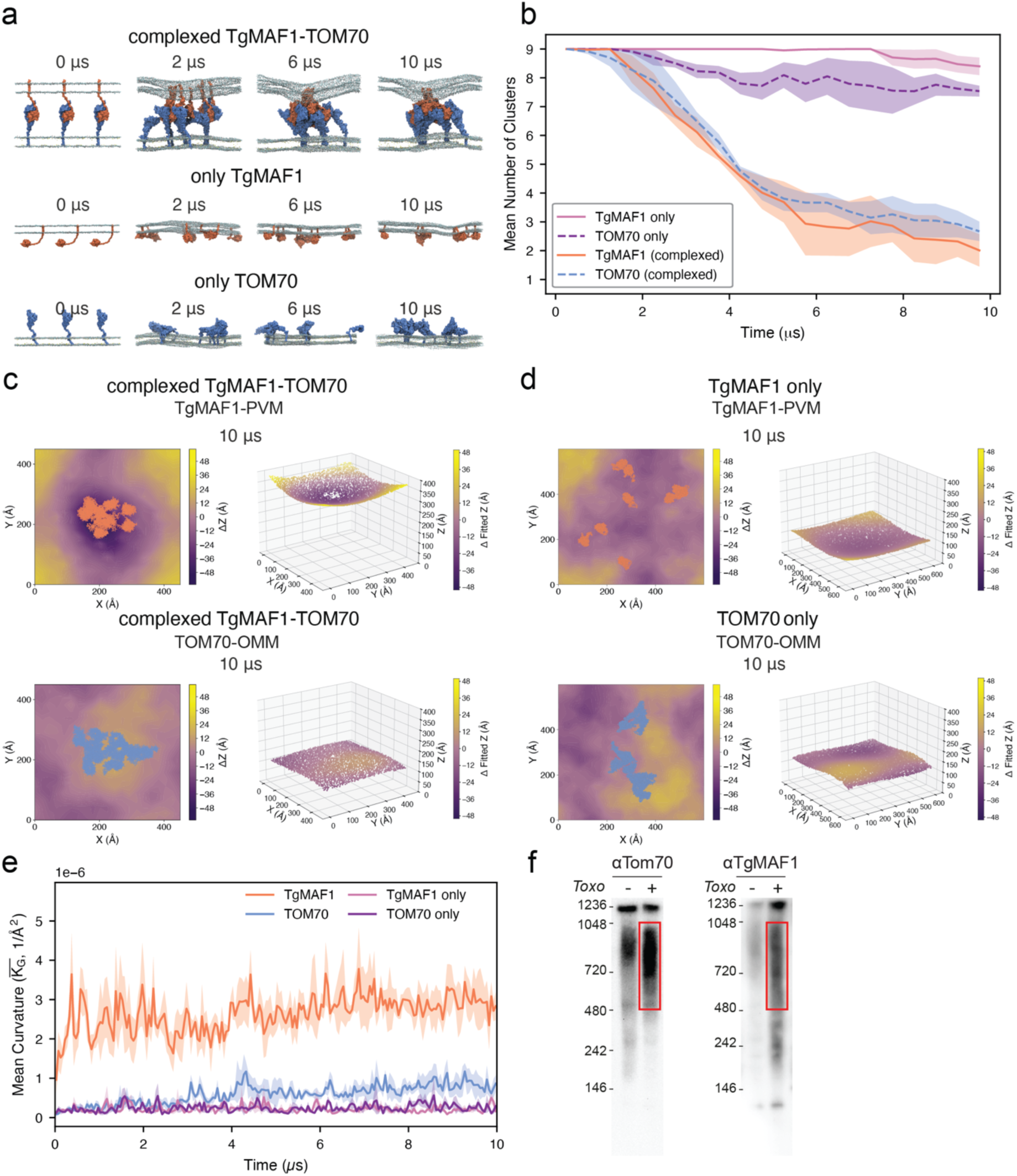
TgMAF1 and TOM70 assemble into higher order complexes that induce local membrane curvature. **a**, (top) Simulation snapshots of complexed or individual TgMAF1-TOM70 of first repeat at 2 μs intervals. 1-palmitoyl-2-oleoylphosphatidylcholine (POPC, grey), 1-palmitoyl-2-oleoylphosphatidylethanolamine and POPE (light blue), and cholesterol (ROH, yellow); bottom, as in top, for TgMAF1 only and TOM70 only systems. **b,** Clustering analysis of TOM70 and TgMAF1 TMH regions following cluster formation over simulation time. Mean cluster number, averaged within 500 ns time bins, is plotted as solid (TgMAF1) and dotted lines (TOM70) with SEM displayed as shaded regions (n=3). **c,** (left) Instantaneous membrane curvature 2D heatmap plots of the PVM and OMM lower leaflets with instantaneous protein coordinates of TgMAF1 (coral) and TOM70 (blue) for complexed TgMAF1-TOM70 system at 10 μs. The deviation from the mean Z height for the PO_4_ head groups of the lower leaflet were plotted to represent the extent of membrane curvature in the simulation, (right) 3D visualization of PO4 head group coordinates fitted to the membrane curvature function and colored by deviation from mean fitted Z coordinates to indicate curvature of PVM and OMM interface facing leaflets for the same system at 10 μs. **d,** (left), as in c (left), for only TgMAF1 or only TOM70 repeat and (right) as in c (right) only TgMAF1 or only TOM70. **e**, Membrane curvature plot of the unsigned mean Gaussian curvature over simulation time within 50 ns time bins for the complexed TgMAF1-TOM70, TgMAF1 only and TOM70 only systems (n=3). Lighter shaded region indicates SEM. **f,** CN-PAGE of solubilized mitochondria from uninfected and infected cells and immunoblotted for TOM70 and TgMAF1, numbers refer to kDa. Data depicted from Repeat 1, Repeat 2 and 3 in Ext. Data Fig. 13-14.

**Table 1.** Cryo-EM data collection, refinement, and modeling statistics for HsTOM70-T*g*MAF1 consensus refinement and local refinement on *Tg*MAF1.

|  | **HsTOM70-TgMAF1<br>composite** | **TgMAF1<br>local refinement** | **HsTOM70-TgMAF1<br>consensus** |
|---|---|---|---|
| PDB ID | 9P0I | - | - |
| EMDB ID | EMD-71074 | EMD-71071 | EMD-71070 |
| **Data Collection** |  |  |  |
| Microscope |  | Talos Arctica | Talos Arctica |
| Camera |  | Falcon 4i | Falcon 4i |
| Magnification (nominal) |  | 190,000 | 190,000 |
| Voltage (keV) |  | 200 | 200 |
| Data acquisition software |  | EPU | EPU |
| Exposure navigation |  | image shift 8 µm | image shift 8 µm |
| Electron exposure (e<sup>-</sup>Å<sup>-2</sup>) |  | 50 | 50 |
| Exposure rate (e<sup>-</sup>/px/s) |  | 7.0 | 7.0 |
| Frames per movie |  | 40 | 40 |
| Pixel size (Å/px) |  | 0.74 | 0.74 |
| Defocus range (µm) |  | -2.0 to -0.8 | -2.0 to -0.8 |
| Recorded movies |  | 13,345 | 13,345 |
| **Image Processing** |  |  |  |
| Processing software |  | cryoSPARC (v.4.6) | cryoSPARC (v.4.6) |
| Final particle images |  | 403,769 | 403,769 |
| Symmetry imposed |  | C1 | C1 |
| Resolution (global) (Å) |  |  |  |
| FSC 0.5 (unmasked/masked) |  | 4.0/4.0 | 4.0/4.0 |
| FSC 0.143 (unmasked/masked) |  | 3.5/3.4 | 3.5/3.4 |
| Local Resolution Range |  | 2.9-4.2 | 2.9-7.6 |
| cFAR |  | 0.45 | 0.51 |
| Map sharpening B-factor (Å<sup>2</sup>) |  | -177 | -122 |
| **Model Refinement** |  |  |  |
| Initial model | AlphaFold3 |  |  |
| Refinement | Coot / ISOLDE<br>Phenix |  |  |
| FSCmap-to-model(0.5) (Å) | 3.6 |  |  |
| MolProbity score | 1.74 |  |  |
| Clash Score | 8.47 |  |  |
| Composition |  |  |  |
| Chains | 2 |  |  |
| Atoms | 5,743 |  |  |
| Protein residues | 733 |  |  |
| Ligands | 0 |  |  |
| Bonds (R.M.S.D.) |  |  |  |
| Length (Å) | 0.004 |  |  |
| Angles (°) | 0.677 |  |  |
| B-factors (min/max/mean) |  |  |  |
| Protein residues | 37.30/151.29/71.00 |  |  |
| Ligand | NA |  |  |
| Ramachandran plot (%) |  |  |  |
| Favored | 95.85 |  |  |
| Allowed | 4.15 |  |  |
| Outliers | 0.00 |  |  |
| Rotamer outliers (%) | 0.00 |  |  |
| Q-score | 0.49 |  |  |
| EMRinger score | 1.40 |  |  |

**Table 2:** Summary of the residue ranges and software used to model the protein structures.

| No. | System | Protein | Modelling | Residue range of final structure |
|---|---|---|---|---|
| 1. | 3x3 TgMAF1 control | TgMAF1 (UniProt ID: A0A140H546) | TMH (93-121aa) + soluble domain (172-446 aa) modelled in ColabFold. Assembled with disordered linker (122-171 aa) in MODELLER. | 92-446 aa |
| 2. | 3x3 TOM70 control | TOM70 (UniProt ID: O94826) | TMH (38-73aa) + soluble domain (114-608 aa) modelled in ColabFold. Assembled with disordered linker (74-113 aa) in MODELLER. | 34-608 aa |
| 3. | 3x3 TgMAF-TOM70 | TgMAF1 (UniProt ID: A0A140H546) | TMH (85-120 aa) + soluble domain (173-446 aa) modelled in PyMOL and AF3 respectively. Assembled with disordered linker (121-172 aa) in MODELLER. | 82-446 aa |
| 4. | 3x3 TgMAF1-TOM70 | TOM70 (UniProt ID: O94826) | TMH (37-60 aa) + soluble domain (109-608 aa) modelled in PyMOL and AF3 respectively. Assembled with disordered linker (61-108 aa) in MODELLER. | 34-608 aa |

**Table 3:** Summary of the membrane composition, box dimensions, length and repeats of coarse-grained molecular dyanmics simulations performed.

| No. | System | PVM Composition* | OMM Composition** | Box size (X x Y x Z) | Length | Repeats |
|---|---|---|---|---|---|---|
| 1. | 3x3 TgMAF1 control | 50% POPC and 50% POPE | - | 61.9nm x 61.9nm x 25.8nm | 10 µs | 3 |
| 2. | 3x3 TOM70 control | - | 50% POPC, 45% POPE and 5% CHOL | 60.8nm x 60.8nm x 33.3nm | 10 µs | 3 |
| 3. | 3x3 TgMAF1-TOM70 | 50% POPC and 50% POPE | 50% POPC, 45% POPE and 5% CHOL | 45.5nm x 45.5 nm x 48.9 nm | 10 µs | 3 |

\*PVM composition adapted from <sup>44, 45</sup>.
\*\*OMM composition adapted from <sup>46</sup>.

This higher-order TgMAF1-TOM70 assembly was accompanied by local membrane bending, as indicated by the marked deviation in PO_4_ headgroup Z-height from the membrane mean plane in both the PVM and OMM interfacial leaflets (**Fig. 4c; Extended Data Fig. 14**). By contrast, quantification of the Gaussian curvature over simulation time confirmed that TgMAF1 or TOM70 alone did not induce membrane curvature (**Fig. 4d-e**; **Extended Data Fig. 14**). To ask whether comparable TgMAF1-TOM70 assemblies form during infection, we performed clear native gel electrophoresis on mitochondria isolated from *Toxoplasma-*infected and uninfected cells. TOM70 was more abundant in 480-1,048 kDa high molecular weight complexes (HMWCs) upon infection, and these comigrated with TgMAF1-containing HMWCs (**Fig. 4f**). Together, these results suggest that TgMAF1 binding to TOM70 induces higher-order complex assembly, and that this assembly drives curvature of the OMM.

## Discussion

This work uncovers a previously unrecognized mode of host subversion in which a parasite effector co-opts the HSP90 chaperone recognition mechanism to target the mitochondrial import receptor TOM70, thereby remodeling a membrane compartment critical for innate immunity and mitochondrial protein import.

TOM70 and HSP90 uniquely occupy fundamental roles in both mitochondrial protein import and innate immune signaling. As an import receptor, TOM70 relies on its TPR clamp domain to dock HSP70/90-bound preproteins at the mitochondrial surface and mediate their translocation across the OMM (Young, Hoogenraad, and Hartl 2003; Scheufler et al. 2000). During viral infection, TOM70 recruits HSP90 bound to IRF3 and TBK1 to the MAVS signalosome to activate antiviral immunity ^34^. This function is targeted by the SARS-CoV-2 protein Orf9b, which upon binding the preprotein-binding groove, impairs HSP90 binding and suppresses interferon signaling (Bouhaddou et al. 2023; Liu et al. 2010). Our findings reveal a distinct mode of TOM70 exploitation: TgMAF1 directly targets the chaperone-binding clamp domain to co-opt TOM70 and induce OMM remodeling. That two evolutionarily distant pathogens converge on TOM70 through distinct mechanisms underscores the importance of this receptor in host cell defense and suggests its manipulation may represent a more general strategy among intracellular pathogens. Indeed, TOM70 interactions have also been predicted for effectors from *Chlamydia,* KSHV, HPV and Dengue virus ^35–38^.

The structural mimicry of HSP70/90 by TgMAF1 is particularly striking. Both the HSP70/90 EEVD motif and the TgMAF1 C-terminal MLD motif engage the same positively charged residues within the TOM70 clamp — K188, R192, and N158 — through analogous electrostatic interactions. This convergence is not coincidental: phylogenetic analyses reveal that the acquisition of a negatively charged, hydrophobic C-terminus in TgMAF1b coincided with the loss of the basic residues present in the ancestral TgMAF1a paralog, consistent with evolutionary selection for HSP90 mimicry. The degree of structural mimicry achieved by TgMAF1 is remarkable given that the overall sequence and structural context of the TgMAF1 C-terminus bears little resemblance to HSP70/90 beyond the terminal residues. This parallels other instances of molecular mimicry in host-pathogen interactions, such as the exploitation of ubiquitin-binding domains by bacterial effectors, in which functional convergence is achieved through minimal sequence elements embedded within structurally unrelated scaffolds ^39^.

Notably, TgMAF1 and HSP90 compete for TOM70 occupancy, raising the possibility that TgMAF1 sequestration of TOM70 and OMM remodeling not only impairs protein import as previously shown, but also impairs antiviral signaling—a potential consequence of infection that warrants future investigation ^15^. The possibility remains that TgMAF1 acts as a functional chaperone mimic that co-opts TOM70 import receptor activity. Two observations are consistent with this idea. First, TgMAF1 traps TOM70 in the open conformation that has an accessible binding cleft, and thus positions TOM70 to interact with mitochondrial preproteins. Second, TOM70 appeared to accommodate both TgMAF1 and Orf9b, suggesting that TgMAF1-bound TOM70 remains competent for substrate binding. Together, these findings raise the possibility that, as a chaperone mimic, TgMAF1 docks at the clamp domain and leaves TOM70 competent for preprotein binding.

Mitochondrial membrane remodeling is canonically driven by dynamin-related GTPases including DRP1 for fission and MFN1/2 for fusion, whose activities are tightly regulated during infection to calibrate immune output (Pernas and Scorrano 2016; Delgado and Pernas 2024). Our findings reveal an orthogonal, GTPase-independent mechanism in which a parasite effector exploits receptor crowding to physically deform the OMM. This is conceptually analogous to the curvature-inducing mechanisms described for BAR domain proteins and reticulon family members, which remodel membrane topology through scaffolding and crowding rather than enzymatic activity ^40–42^. The mechanism we describe has broader implications for understanding how intracellular pathogens manipulate membrane architecture, although whether TgMAF1-induced OMM curvature requires additional host factors remains an important open question.

Finally, our results illuminate the evolutionary logic of TgMAF1 as a host subversion factor. The neofunctionalization of TgMAF1b from an ancestral paralog that lacks TOM70 binding activity by the selective acquisition of an HSP90-like C-terminal motif is representative of molecular mimicry through effector evolution. This parallels the HopBF1 kinase of the plant pathogen *P. syringae*, which mimics an HSP90 client to phosphorylate the HSP90 active site, thereby inactivating a central node of plant immunity ^43^.

In summary, we have identified TgMAF1 as a chaperone mimic that engages the HSP70/90-binding clamp of TOM70 to nucleate higher-order protein assemblies and drive local OMM curvature during *Toxoplasma* infection. Our findings establish molecular mimicry of chaperone recognition motifs as a strategy for pathogen-induced membrane remodeling, and position TOM70 as a convergent target of diverse intracellular pathogens. Future work defining the downstream consequences of TOM70 sequestration for mitochondrial protein import and innate immune signaling will be essential to understand the consequences of *Toxoplasma* exploitation of mitochondrial architecture during infection.

## Acknowledgments

We thank the MPI-AGE FACS and Imaging Core for microscopy support; the CECAD electron microscopy facility, in particular Katrin Seidel. We thank William Lessin at The Scripps Research Institute Hazen cryo-electron microscopy facility for microscope support. We thank all members of the Pernas laboratory for helpful discussions. We acknowledge VEuPathDB as an essential resource for this work and thank all its contributors. Molecular graphics and analyses performed with UCSF ChimeraX, developed by the Resource for Biocomputing, Visualization, and Informatics at the University of California, San Francisco, with support from National Institutes of Health R01-GM129325 and the Office of Cyber Infrastructure and Computational Biology, National Institute of Allergy and Infectious Diseases.

## Author contributions

Conceptualization: J.S., S.K. and L.P. Methodology: all authors. Investigation: All authors. Resources: All authors. Funding acquisition: L.P. Project administration: L.P. Writing, original draft: L.P. Review and editing: All authors. Supervision: L.P.

## Data and materials availability

All density maps and models reported in this article were deposited in the Electron Microscopy Data Bank (EMDB) and the Protein Data Bank (PDB). The consensus reconstruction can be accessed EMD-71070. The local refinement for MAF1 was deposited as EMD-71071. The composite is available as EMD-71074 and PDB-9P0I. Molecular dynamics simulation coordinate and trajectory files will be deposited on Zenodo.

## This file contains

Figures 1 to 4

Extended Data Figures 1 to 14

## Other supplementary materials for this manuscript include the following

Table 1–3

Ref 44-81

## Materials and Methods

### Cell culture and cell lines

HeLa adenocarcinoma cells, Human foreskin fibroblasts (HFF), were obtained from ATCC (CCL-2, and SCRC-1041, respectively); HeLa cells stably expressing pMXs-Cox4-BFP were generated through lentiviral transduction. TOM70-GFP expressing MEFs were obtained from Drs. Janet Shaw and Adam Hughes (U. Utah). TOM70-GFP expressing MEFs were engineered to express TOM70-BFP mutants through lentiviral transduction. Cells were tested every 2 weeks for *Mycoplasma* infection by means of polymerase chain reaction (PCR).

### Parasite culture and generation of parasite strains

All parasite strains were maintained by serial passage on HFF monolayers in cDMEM. *Toxoplasma* strains used in this study include: Type I RH*Δhxgprt*:mCherry+ (*Toxo*^mCherry^); Type I RHΔ*maf1:mCherry+* (Δ*maf1*); RHΔ*maf1:HA-MAF1:mCherry+* (Δ*maf1*:HA-MAF1); and ^13^, ^15, 47^. All strains were routinely tested for *Mycoplasma* infection by polymerase chain reaction (PCR) every two weeks. The TgMAF1 mutants: Δ*maf1*:HA-MAF1^D446Δ^; Δ*maf1*:HA-MAF1^D446E^; Δ*maf1*:HA-MAF1^D446N^; Δ*maf1*:HA-MAF1^ML444QR^; Δ*maf1*:HA-MAF1^ML444QG^; and Δ*maf1*:HA-MAF1a^QR444QG^ were generated via site directed mutagenesis from either the Δ*maf1*:HA-MAF1 or Δ*maf1*:HA-MAF1a plasmids. To create transgenic parasites, RHΔ*maf1:mCherry+* were transfected with 60 µg of the plasmids harboring the mutations following NotI linearization and selected with 20 µM chloramphenicol (Sigma: R4408). 2 weeks post-selection, the populations were cloned out via serial dilution. Single clones were confirmed with HA staining in immunofluorescence assays.

### Vector generation

Site-directed mutagenesis was performed using the following primers:

F_MAF1^D446E^ (TAGCATGCTGgagTGAGCGGCCG);

R_MAF1^D446E^ (GCCAGATACGTGCTTTCAGCCTC);

F_MAF1^D446Δ^ (TGAGCGGCCGCTACGTAA);

R_MAF1^D446Δ^ (CAGCATGCTAGCCAGATACG);

F_MAF1^D446N^ (TAGCATGCTGaacTGATTAATTAAG);

R_MAF1^D446N^ (GCCAGATACGTGCTTTC);

F_MAF1^ML444QG^ (TCTGGCTAGCcagggaGACTGATTAATTAAGACTACGACGAAAG);

R_MAF1^ML444QG^ (TACGTGCTTTCAGCCTCCTGT) F_MAF1^ML444QR^ (TCTGGCTAGCcagagaGACTGATTAATTAAGACTACGACG);

R_MAF1^ML444QR^ (TACGTGCTTTCAGCCTCCTG);

F_MAF1a^QR444ML^ (CAAGTCTTATatgctaGACTGATTAATTAAGACTAC);

R_MAF1a^QR444ML^ (AACTTCCTTTCAGATTCCAG);

### Alphafold modeling

Protein sequences were retrieved from UniProt and modeled using AlphaFold2 without template constraints using our in-house pipeline. For each input, 25 models were generated; the top-ranked model (rank 0) based on pLDDT was selected. The predicted aligned error (PAE) matrix was used to assess residue-pair confidence. For TgMAF1b (Uniprot: A0A140H546), the unstructured N-terminus up to and including the proline-rich region were excluded (residues 1-173 were deleted and residues 174-446 were included). TgMAF1a (Uniprot: A0A7J6KEI3) was truncated to the same length for comparison (residues 1-162 were deleted and residues 163-435 were included). TOM70’s (Uniprot: O94826) transmembrane domain was also removed due to disordered structure (residues 1-111 were deleted and residues 112-608 were included). PAE plots were generated from AlphaFold2 output pkl files. The protein 3D structures were then visualized using the PyMOL software (The PyMOL Molecular Graphics System, Version 3.1.4 Schrödinger, LLC). For modeling protein complexes using AlphaFold3 (AF3), sequences were likewise retrieved from UniProt and modeled using the AlphaFold server. For each input, 5 models were generated; the top-ranked model (rank 0) was selected. The AF3 predicted aligned error (PAE) matrix was used to assess residue-pair confidence. The cytosolic domain of HsTOM70 (Uniprot: O94826; amino acids 111-608) was modeled together with TgMAF1b (Uniprot: A0A140H546; amino acids: 173-446) and a minimal peptide of SARS-CoV-2 Orf9b (Uniprot: P0DTD2, amino acids 44-70). PAE plots were generated by AlphaFold3. The protein 3D structures were then visualized using the ChimeraX software.

### Electron microscopy sample preparation and analysis

HeLa cells were grown on small discs of aclar foil (Science Services, #E50425-10) in 12-well plates and infected with *Toxoplasma*. 5 hours post infection (hpi), discs were fixed for 1 h in 2% glutaraldehyde (Sigma: G5882-100ML) with 2.5 % sucrose (Roth: 4621.1) and 3mM CaCL_2_ (Sigma, # C7902-500G) in 0.1M HEPES buffer (Sigma: C7902-500G) pH 7,4. Samples were washed three times with 0.1M HEPES buffer and incubated with 1% osmium tetroxide (Science Services: E19190) and 1% potassium hexacyanoferrate (Sigma: P8131) for 1 h at 4°C. After 3×5 min wash with 0.1M cacodylate buffer (Applichem: A2140,0100), samples were dehydrated at 4°C using ascending ethanol series (50%, 70%, 90%, 3×100%) for 7 min each. Infiltration was performed with a mixture of 50% Epon/ethanol for 1h, 70% Epon/ethanol for 2h and with pure Epon (Sigma: 45359-1EA-F) overnight at 4°C. Samples were embedded into TAAB capsules (Agar Scientific: G3744) and cured for 48 h at 60°C. Ultrathin sections of 70 nm were cut using an ultramicrotome (Leica Microsystems, UC6) and a diamond knife (Diatome, Biel, Switzerland) and stained with 1.5 % uranyl acetate (Agar Scientific: R1260A) for 15 min at 37°C and 3 % Reynolds lead citrate solution made from Lead (II) nitrate (Roth: HN32.1) and tri-sodium citrate dehydrate (Roth: 4088.3) for 4 min. Images were acquired using a JEM-2100 Plus Transmission Electron Microscope (JEOL) operating at 80kV equipped with a OneView 4K camera (Gatan). To measure the percentage of the PVM associated with host mitochondria, the total length of contacts between the mitochondria was totaled and divided by the perimeter of the PVM (PVM associated with mitochondria /total PVM perimenter ×100) using ImageJ.

### Immunofluorescence assays and antibodies

TOM70-GFP expressing MEFs were plated in a 24 well glass-bottom plate (Greiner Bio-One), infected with *Toxoplasma* strains and harvested at 5 hpi. Plates were fixed in 4% paraformaldehyde in prewarmed cDMEM for 15 min at 37°C, permeabilized for 10 min at RT with 0.2% triton, blocked in 3% BSA for 30 min, incubated in 1:250 of primary Ab O/N at 4°C. After 3×5 min washes with 1X PBS, cells were incubated in 1:1000 anti-mouse Alexa Fluor Plus for 40 min to 1 h at RT. Plates were rinsed 3X in 1X PBS and maintained in 1X PBS until imaging. Primary Abs: TOMM70 (HPA:048020); Antisera of TgMAF1^13^; HA (CST:3724, Roche 3F10); Secondary Abs: Alexa Fluor Plus 405, Alexa Fluor Plus 488, Alexa Fluor Plus 594, Alexa Fluor Plus 647 (Thermo Fisher). Images were taken with a 60X/1.35 or 100X/1.35 silicon oil objective and excitation with either 405, 488, 561, or 640 confocal or Olympus super resolution laser lines with an Olympus IXplore SpinSR spinning disk confocal microscope.

### Immunoprecipitation/Immunoblotting

MEF cells or HeLa cells were seeded at either 5 million in 15 cm dishes or at 2.2 million in 10 cm dishes. The next day, cells were infected with *Toxoplasma* at a multiplicity of infection of 6 and harvested 24 hours post infection. Infected cells were scraped down in 1XPBS (Gibco) containing protease (Thermo Scientific) and phosphatase (Roche) inhibitors and incubated in lysis buffer for 15 min at 4°C. Cleared lysates were incubated with 25 μl magnetic HA-beads (Thermo Scientific: 88837) or 25 μl magnetic GFP-nanobodies (Chromotek GTD-20) overnight. The beads were washed 3x times with 1XPBS (Gibco) containing protease (Thermo Scientific) and phosphatase (Roche) inhibitors. Afterwards, the samples were eluted from the magnetic beads with 2X SDS buffer by incubating them at 40°C for 10 min. Samples were processed for gel electrophoresis and probed with indicated antibodies. For immunoprecipitation in the presence of HSP90 peptides, cleared lysates were prepared as described above. The peptides Ac-DDTSRMEEVD or Ac-DDTSRMEEVN were diluted in nuclease-free water to a final volume of 100 μl and added to the lysates to reach the desired final concentration. The samples were then eluted with 2X SDS loading buffer from the magnetic beads by incubating them at 40°C for 10 min. At the end 10 μl input (1%) and 20 μl elution (23%) were electrophorized using 10% or 4-20% SDS-PAGEs (Invitrogen) and transferred to a PVDF membrane (Immobilion-P). Using a ECL Chemostar scanner (Intas) the membrane was scanned for probed antibodies. GFP (Takara Bio: 632380); TOM40 (Sigma-Aldrich: HPA036231); VDAC2 (Proteintech; 11663-1-A); HA-HRP (Roche 12013819001); TgMAF1b polyclonal generated in ^48^ were used at 1:300 or 1:2000 O/N.

### Live cell imaging

Cells were seeded on 24-well CELL view glass bottom cell culture plates (Greiner Bio-One) and imaged using an Olympus IXplore SpinSR 50 mm spinning disk confocal microscope. Live cell imaging was performed with incubation at 37°C and 5% CO_2_. Images were taken with a 100X/1.35 silicon oil objective and excitation with either 488, 561, or 640 laser lines and cellSens Software.

### Immunoblotting and antibodies

Uninfected or *Toxoplasma*-infected cells were harvested in chilled lysis buffer [50mM Hepes-KOH pH 7.4, 40mM NaCl, 2mM EDTA, 1% Triton X-100 and protease and phosphatase inhibitors (Thermo Scientific: A32961 and Sigma: 4906845001) (lysis buffer) and lysed for 30 min on ice. Lysates were then centrifuged at 10 minutes at 14,000 x g at 4 °C and the clarified supernatant was transferred to a fresh tube with 5X SDS for a final volume of 1X SDS. Protein lysates were resolved in SDS-PAGE and following gel transfer, the membranes were blocked with TBS-0.05% Tween 20 (TBS-T) and 5% milk and the primary antibodies were incubated overnight. Following incubation, blots were washed three times in TBS-T for 15 minutes and then incubated with horseradish peroxidase (HRP)-conjugated anti-rabbit IgG (CST: 7074) or anti-mouse IgG (CST: 7076) at a 1:4000-1:10,000 dilution for ∼1 h and developed using a chemiluminescence system (Pierce™ ECL Western Blotting Substrate or Pierce SuperSignal™ West Atto Ultimate Sensitivity Substrate; ThermoFisher Scientific). The following primary antibodies were used: TOMM70 (HPA048020); HA-HRP (Roche 12013819001 TOMM40 (HPA036231); VDAC2 (Proteintech 11663-1-A)

### Peptide synthesis

The peptides Ac-DDTSRMEEVD-NH2 (MW_calc_ = 1238.23 Da; MW_exp_ = 1238.28, final purity ≥ 98 %) and Ac-DDTSRMEEVN-NH2 (MW_calc_ = 1237.24 Da; MW_exp_ = 1238.08, final purity ≥ 98 %) were synthesized by solid-phase peptide synthesis on a peptide synthesizer (Syro I, MultiSynTech) using the fluorenylmethoxycarbonyl (Fmoc)/*tert*-butyl strategy on a Rink amide resin (0.48 mmol/g, 15 μmol scale). The amino acid coupling steps were performed twice for each amino acid using eight equivalents each of OxymaPure (2-cyano-2(hydroxyamino)acetate), DIC (dicyclohexylcarbodiimide), and the respective Fmoc-protected amino acid in DMF (dimethylformamide). The protecting group was removed by incubating the resin first in 40% piperidine in DMF followed by 20% piperidine in DMF. All peptides were N-terminally acetylated by shaking them for 15 min at RT in an acetic anhydride/ DIPEA (Diisopropylethylamine)/ DCM (dichloromethane) (75:12.5:12.5, v/v/v) mixture. Peptides were cleaved from the resin using a mixture of trifluoroacetic acid/ thioanisole/ 1, 2-ethanedithiol (90:7:3, v/v/v). The crude peptides were purified by reverse-phase HPLC using a linear gradient of 10– or 20–60% B in A (A: water/0.1 % TFA; B: acetonitrile (ACN)/0.1 % TFA) over 45 min.

### Mitotracker labeling

Medium was replaced on uninfected or infected cells with prewarmed cDMEM containing MitoTracker Deep Red at a concentration of 25 nM. After 20 min of incubation at 37°C, media was replaced prewarmed cDMEM.

### Bioinformatics

Sequence conservation in Fig. 3c was performed using WebLogo 3 ^49, 50^. For Supp. Fig. 9, T-COFFEE Simple Multiple Sequence Alignment was used to generate the sequence alignments ^51, 52^.

### Expression and purification of TgMAF1-TOM70 complexes

*T. gondii* RH MAF1b (residues 173-446, TgMAF1) and human TOM70 (residues 111-608) were co-expressed using a pRSF-Duet vector backbone where TOM70 was N-terminally tagged with a 6x polyhistidine (6xHis) tag and TgMAF1 was N-terminally tagged with a StrepII tag. *E. coli* BL21(DE3) were transformed with the above-mentioned vector and grown in LB_Kan_ (1% w/v peptone, 0.5% w/v yeast extract, 1% w/v NaCl, 50 ug/ml Kan) at 37 °C to an OD_600_ of 0.8. Expression was induced by addition of 1 µM IPTG and proteins were expressed for 4 h at 37 °C. Bacteria were harvested at 5,000 xg for 20 min at RT, washed with PBS and frozen until further use. For small batch purifications frozen cell pellets were resuspended in 2.5 ml cold BugBuster Master Mix (Millipore Sigma) per gram pellet supplemented with protease inhibitor cocktail (Thermo Scientific) and 2 µM PMSF and lysed at 4 °C for 5 hours on a rolling shaker. The lysate was cleared by centrifugation at 21,000 xg for 15 min at 4 °C and 1 ml supernatant was added to 400 µl MagStrep Strep-Tactin XT (IBA, Cat # 2-5090-002) bead slurry equilibrated with Buffer W (IBA, Cat # 2-1003-100) and incubated over night at 4 °C on a rotary mixer. The beads were washed 4 times with buffer W, transferred to a fresh reaction tube and proteins were eluted with 200 µl BXT buffer (IBA, Cat # 2-1042-025) for 20 min at 4 °C on a thermomixer at 800 rpm. For larger preparations of isolated TgMAF1-TOM70, frozen pellets were resuspended in 3 ml wash buffer (100 mM Tris, 150 mM NaCl, 2 mM EDTA) supplemented with 1 mM PMSF, benzonase, and 0.1% LDAO per gram pellet and cells were lysed using an Emulsiflex-C3 homogenizer (Avestin). The lysate was cleared by centrifugation at 35,000 xg for 25 min and loaded onto a 5 ml StrepTactin XT column (IBA, Cat # 2-5010-002) at 1 ml/min. Proteins were eluted with wash buffer supplemented with 50 mM biotin. Fractions containing TgMAF1-TOM70 complex were pooled and subjected to size exclusion chromatography for further purification using a Superdex 200 increase 10/300 GL column (Cytiva) in wash buffer at a flow rate of 1 ml/min. Peak fractions were collected and concentrated using an Amicon Ultra-15 30 kDa concentrator. Protein samples were snap-frozen and stored at 80 °C until further use. The molecular weight of the isolated TgMAF1-TOM70 complex was estimated by comparison with a gel filtration standard (Biorad, Cat # 1511901).

### Clear-native (CN) PAGE of TgMAF1-TOM70 from mitochondrial fractions

For the isolation of TgMAF1-TOM70 from mitochondria of infected cells, 5 M HeLa cells were seeded into 15 cm dishes 36 hours prior to infection. *T. gondii* RH were freshly lysed by trituration from at 24 hours post infection (hpi) from human foreskin fibroblast (HFF), isolated by centrifugation at 550 xg for 5 min at 37 °C, resuspended in fresh media and filtered through a 5 µM syringe filter. HeLa cells were infected at a multiplicity of infection (MOI) of 10 and at 2 hpi, the infection was synchronized by removing the media containing uninvaded parasites through aspiration and replacing with fresh media. At 18 hpi, cells were washed with 10 ml cold PBS, harvested by scraping in 1.5 ml MS buffer (225 mM mannitol, 75 mM sucrose, 30 mM Tris/Cl, 1 mM EDTA, pH 7.4), and lysed by 10 passages through a 27G needle on ice. The nuclear fraction was removed by centrifugation at 700 xg for 5 min at 4 °C. To improve mitochondrial yield, the nuclear fraction as overlayed with 500 µl MS buffer and again triturated by 10 strokes through a 27G needle. Mitochondria were isolated from the combined post-nuclear fraction by centrifugation at 10,000 xg for 15 min at 4 °C. The cytosolic fraction was removed, and mitochondria were washed two times with 1 ml MS buffer. After the final centrifugation, mitochondria were resuspended in 200 µl MS buffer per 15 cm dish HeLa cells and total protein concentration was quantified by BCA assay. For clear-native polyacrylamide gel electrophoresis (CN-PAGE), 100 ug mitochondria were sedimented for 10 min at 10,000 xg at 4 °C, resuspended in 11.5 µl Sol A buffer (50 mM NaCl, 50 mM imidazole/HCl, 2 mM 6-aminohexanoic acid, 1 mM EDTA, pH 7.0) containing 1% digitonin (Millipore Sigma, Cat # 300410) and incubated for 15 min on ice. Samples were cleared by centrifugation at 20,000 xg for 20 min at 4 °C, supplemented with CN loading buffer (50% glycerol, 0.1% Ponceau) and loaded onto a 4-16% native gel (Thermo Scientific, Cat # BN1004). Gels were run in native PAGE running buffer (Thermo Scientific, Cat # BN2001) at 80 V for 4-6 h at 4 °C. The molecular weight of sample proteins was estimated by comparison with the NativeMark protein standard (Thermo Scientific, Cat # LC0725). Gels were subsequently stained with SimplyBlue (Thermo Scientific, Cat # LC6060) for 1 h at RT and destained for 2 h in water. Destained gels were incubated in cold transfer buffer for 15 min before Western blotting for 2 h at 200 mA in transfer buffer (10 mM CAPS (3-(cyclohexylamino)-1-propane sulfonic acid), 10% (v/v) EtOH, pH 11) at 4 °C. Membranes were subsequently incubated for 15 min in 8% acetic acid and washed 2x in water and once in ddH2O before air-drying the membranes at RT for 30 min. Dried membranes were activated in MeOH for 5 min at RT, washed once with water, 2x with TBS-T (20 mM Tris/Cl, 150 mM NaCl, 0.05% Tween 20, pH 7.6), and blocked with 5% BSA-TBS for 30 min at RT. Primary antibodies (TOM70: Millipore Sigma, Cat # HPA014589, 1:1000; MAF1: homemade, 1:1000) were incubated in 5% BSA-TBS over night at 4 C, membranes were washed 3x in TBS and once in TBS-T before secondary antibody decoration in 5% milk-powder-TBS for 1 h at RT. Membranes were washed 4x with TBS-T and proteins were detected using ECL Western blotting substrate (Thermo Scientific, Cat # 32106).

### Yeast strains, cultivation and mitochondria isolation

The bakeŕs yeast strains YPH499 and tom70Δ have been described (Becker et al., 2011). pRS416 plasmid expressing *TOM70* was used to generate *TOM70 N137A* and *TOM70 N174A* by site-directed mutagenesis. Subsequently, these constructs were transformed into the tom70Δ strain to obtain yeast expressing *TOM70 N137A* or *TOM70 N174A*. The strains were grown in selective medium (SC-Ura + 2 % glucose) at 30°C. Yeast cells were grown until early logarithmic growth phase. Mitochondria were isolated by differential centrifugation^53^. In brief, yeast cells were harvested by centrifugation (5,500 g, 5 min, 4°C) and washed with water. After treatment with DTT buffer (10 mM DTT, 100 mM Tris/HCl pH 9.4), cells were incubated with 4 mg zymolyase per g of cells in zymoylase buffer (1.2 M sorbitol, 20 mM KPi pH 7,4) for 45 min at 30°C. The cell wall was ruptured by homogenization using a glass potter in homogenization buffer (0.6 M sorbitol, 10 mM Tris/HCl pH 7.4, 1 mM ethylenediaminetetraacetic acid (EDTA), 1 mM phenylmethylsulfonylfluoride (PMSF), 0.2% (w/v) bovine serum albumin). Cell debris and fragments were removed by centrifugation. Subsequently, mitochondria were isolated by centrifugation (17,000 g, 15 min, 4°C) and washed with SEM buffer (250 mM sucrose, 10 mM MOPS/KOH pH 7.2, 1 mM EDTA). Aliquots with protein concentrations of 10 mg/ml were shock-frozen in liquid nitrogen and stored until use.

### Import assay

^35^S-labelled Aac2 was produced by in vitro translation using a cell-free translation extract (TNT® Quick Coupled Transcription/Translation System, Promega) and pGEM4z encoded Aac2 as template^53^. The import reaction was performed following an established procedure. The Aac2 lysate was incubated with mitochondria in import buffer (1% (w/v) BSA, 250 mM sucrose, 80 mM KCl, 5 mM MgCl2, 10 mM MOPS/KOH pH 7.2, 5 mM methionine, 2.5 mM KH_2_PO_4_ pH 7.2, 2 mM ATP, 2 mM NADH, 120 μg/ml creatine kinase, 12 mM creatine phosphate) for different time periods at 26°C. The import reaction was stopped by addition of 1 μM valinomycin to deplete the membrane potential. Mitochondria were re-isolated and washed with SEM buffer. Subsequently, mitochondria were resuspended in lysis buffer (20 mM Tris-HCl pH 7.4, 0.1 mM EDTA, 50 mM NaCl, 10% glycerol, 1 mM PMSF) containing 1% (w/v) digitonin for 15 min on ice. After a clarifying spin, the lysate was analyzed by blue native electrophoresis (Priesnitz et al., 2020).

### Negative staining TEM

Sample quality was assessed by negative-stain electron microscopy following established protocols ^54^. In brief, 200 nM of TOM70-MAF1 was applied to glow-discharged 400 square Mesh Copper/Rhodium Maxaform grids (Electron Microscopy Sciences) and stained with 2 % uranyl formate. Micrographs were collected automatically with Leginon on a Thermo Fisher Talos F200C transmission electron microscope operating at 200 keV and equipped with a Gatan K2 direct electron detector at a nominal magnification of 73,000, corresponding to a pixel size of 1.981 Å. Micrographs were imported in cryoSPARC (v.4.6.2) with constant CTFs and inverted contrast. Templates were generated from manually picked particles. 2D classification without Force max over poses/shifts yielded well-aligned 2D averages, suitable for cryo-EM studies (Extended Data Fig. 2).

### Preparation of samples for cryo-EM

Strep-Tactin elutions were thawed on ice and aggregates were pelleted by centrifugation for 10 min at 15,000 g. The resulting 2 mg/ml sample in 100 mM Tris, 150 mM NaCl, 1 mM EDTA, pH 8 was used for cryo-EM. UltraAuFoil 300 mesh R 0.6/1.0 grids were glow-discharged under vacuum for 30 s at 15 mA in a Pelco easiGlow 91000 glow discharge cleaning system (Ted Pella). 3.5 µl sample was applied to the front of the grid and blotted with Whatman 1 filter paper for 4 s after the liquid spot on the filter paper stopped spreading. To overcome preferred orientation, 7.5 mg/ml sample was vitrified with the addition of 0.08% (v/v) decyl-maltoside(n-decyl-β-D-maltopyranoside) and the blotting time was increased to 6 s. Grids were manually plunge-frozen in a 4 °C cold room with >95% humidity. AutoGrids were assembled and immediately screened.

### Cryo-EM data collection and processing

Cryo-EM datasets were collected on a Talos Arctica TEM (Thermo Fisher) operating at 200 keV. Movies were recorded using a Falcon 4i direct electron detector (Thermo Fisher) at a nominal magnification of 190,000, corresponding to a pixel size of 0.74 Å. Movies were saved in the electron-event representation (EER) format and recorded at a total electron exposure of 50 e^−^ per Å^2^. All datasets were collected automatically using EPU (v.3.9, Thermo Fisher) with a defocus range of −0.8 to −2.0 μm. EPUs fast exposure navigation was used to collect data at an 8 µm image shift. All datasets were processed using cryoSPARC (v.4.6). The processing workflow is shown in Extended Data Fig. 3. A total of 13,345 movies were collected and dose-fractioned into 40 frames, motion corrected using patch-based motion correction, followed by patch-based CTF estimation in cryo-SPARC live. Micrographs with CTF fits above 6 Å and astigmatism above 600 Å were discarded. The remaining 11,948 accepted micrographs were used for further processing.

During live data collection, 60-150 Å blob-picked particles were used for 2D classification to generate templates. These particles were extracted in a 320 px box, Fourier cropped to 100 px. Using these templates and a 100 Å particle diameter, eight million particles were picked and subjected to 2D classification. 2D classes with a balanced range of viewing directions were selected to train TOPAZ ^55^. 216 exposures with 100-110 particles per micrograph, corresponding to the selected 2D classes, were used for training, giving 22,645 particles. The training on non-denoised micrographs with pre-train initialization (ResNet16) was done with a 100 Å estimated particle diameter, a learning rate of 0.0001, a minibatch size of 64, 20 epochs, an epoch size of 10,000, and a train-test split of 0.3. TOPAZ extract was run with a radius of extracted regions of 1 and a particle threshold of 6, yielding 4.7 million particles. 4.4 million particles were subjected to 2D classification using 120 classes, a mask diameter of 120 Å, an uncertainty factor of 3, a minimum separation distance of 20 Å to remove duplicates, 40 online-EM iterations, and a batch size per class of 400. Combining the template and TOPAZ picked particles resulted in 1.6 million particles after duplicate removal.

Using the ab initio reconstruction from cryo-SPARC live and two decoy classes, particles were classified using heterogeneous refinement (HR) with a batch size per class of 2,000 and a spherical mask diameter of 110 Å. 882 thousand particles were selected and further refined with non-uniform refinement (NUR) with a 15 Å lowpass-filtered input and re-extracted using the full 320 px box-size. A second round of NU and HR removed additional low-quality particles, resulting in 810K particles. To further improve the resolution, focused 3D classification without alignment was used. A mask was generated around the local MAF1 density, using no dilation and a padding of 15 px. 3D classification ran, using a 4 Å filtered resolution in input mode with 4 classes. Half of the dataset was selected (403 thousand particles) and subjected to a local refinement with recentering using a mask covering the entire complex. Gaussian priors were enabled with search priors of 6° rotation and 2 Å shifts. This gave the final consensus reconstruction with a global resolution of 3.5 Å (FSC 0.143, B-factor - 122 Å^2^). Local refinement with a tight mask on the MAF1 subunit using 12 px padding and the above-mentioned non-default parameters yielded a 3.5 Å reconstruction (FSC 0.143, B-factor - 177 Å^2^). Reported B-factors result from unsupervised auto-sharpening during refinement in cryoSPARC. A full validation of the reconstruction in shown in Extended Data Fig. 4-5.

### Model building, refinement, and validation

To aid model building, unsharpened half-maps, FSC-based resolution, and molecular mass were supplied as input to generate a composite map within Phenix (v.1.19) ^56^. The resulting composite map was further modified using EMReady, which improved interpretability during model building ^57^. The initial atomic models were generated with AlphaFold3, providing the sequences for HsTOM70^111-608^ (O94826) and *Tg*MAF1^173-446^ (A0A140H546) ^28^. The Alphafold model was fitted into the densities as rigid bodies within ChimeraX and manually corrected using Coot (v.0.9.8.95) (https://doi.org/10.1002/pro.3791) and ISOLDE and iteratively refined using *phenix.real_space_refine* within Phenix (v.1.19) ^58^. Validation reports were automatically generated using MolProbity within Phenix ^58^. Q-scores were calculated within ChimeraX (https://github.com/tristanic/chimerax-qscore) ^59^. Local motion trajectories were visualized with our custom python script (https://github.com/schaefer-jh/motionviz). Sequence conservation for TOM70 (O94826) and MAF1 (A0A140H546) was calculated using default parameters on the consurf server (https://consurf.tau.ac.il/). The full validation report is provided in Table 1.

### Molecular Dynamics simulations

TgMAF1b and TOM70 were modelled using a combination of ColabFold, AlphaFold3 (AF3), PyMOL (version 3.0), and MODELLER ^60^ ^28, 61^. The modelling details of the proteins in each system are listed in Table 2. The final models were then converted to coarse-grained (CG) representations using the Martinize2 script and Martini 3 force field prior to building the system ^62, 63^. The default Martinize2 elastic network was applied using a force constant of 700 kJ mol⁻¹ nm⁻² between non-successive backbone beads with lower and upper cutoff of 0.0 nm and 0.9 nm respectively. The protein models were then orientated using PPM 3.0 web server prior to embedding in a membrane bilayer ^64^.

The coarse-grained protein models were embedded into a membrane bilayer using the INSANE script ^65^. Lipid compositions for PVM and OMM using 1-palmitoyl-2-oleoylphosphatidylcholin (POPC) and 1-palmitoyl-2-oleoylphosphatidylethanolamine (POPE) and cholesterol are shown in Table 2. PVM composition adapted from ^44, 45^ and OMM composition adapted from ^66^. An in-house modified version of the INSANE script was used to incorporate the Martini 3 cholesterol model ^67^. The separation between the OMM and PVM was set to ∼20 nm prior to energy minimization and equilibration. All systems were minimized by steepest descents (5000 steps) before solvating with martini CG waters using gmx solvate and neutralizing with 0.15 M of NaCL using gmx genion in GROMACS 2021^68^.

Systems were minimized and equilibrated using the CHARMM-GUI membrane builder protocol files ^69^. The minimization step consisted of a steepest descent integrator over 550,000 steps. The temperature was maintained at 303.15 K using the V-rescale thermostat and the pressure was semi-isotropically maintained at 1.0 bar using Berendsen barostat ^70, 71^. Post minimization, the systems were subjected to five equilibration steps using leap-frog integrator with position restraints with decreasing force constants from 1000 kJ/mol/nm² to 50 kJ/mol/nm² for protein backbone and from 200 kJ/mol/nm² to 10 kJ/mol/nm² for lipid headgroups. Post equilibration, the primary 1×1 systems were replicated using genconf-nbox to create the 3×3 systems. The production run employed a md integrator for a total of 10 μs at 0.02 ps time step. The temperature was maintained at 303.15 K using a V-rescale thermostat that was applied to the protein, membrane and solute groups separately. Pressure was semi-isotropically maintained at 1.0 bar with Parinello-Rahman barostat ^72, 73^. Further, to minimize potential bias in protein clustering, an additional equilibrium stage was performed for second and third repeats for the TgMAF1-TOM70 system. Specifically, a 1 μs position restrain simulation, with 1000 kJ/mol/nm² force constant applied to the first atom of TOM70 was performed prior to the 10 μs production run to allow for rotational movements of the protein complex. Production simulations were run on ARCHER2 using GROMACS 2022.4 (https://www.archer2.ac.uk). Table MD3 summarizes the simulations performed in this study.

Systems were visualized using VMD ^74^. MDAnalysis was used for trajectory manipulation ^75, 76^. For minimum distance analysis (mindist analysis) gmx mindist was used. For clustering analysis Density-Based Spatial Clustering of Applications with Noise (DBSCAN) algorithm within the Scikit-Learn library was used ^77, 78^. An epsilon (ε) value of 20 nm and a minimum sample size of 1 was used to define cluster neighborhood for the protein TMHs with their centers of masses used as reference points. Scripts for the 2D heatmap visualizations of membrane curvature was adapted from Chavent et al., 2015. Membrane curvature analysis methods were adapted from ^79^. Numerical operations, statistical analyses, and data visualization were conducted using NumPy^78, 80–82^.

### Statistical analyses

All statistical analyses were performed using GraphPad Prism 9 software and are indicated accordingly.

---

**Extended Data Fig. 1:**
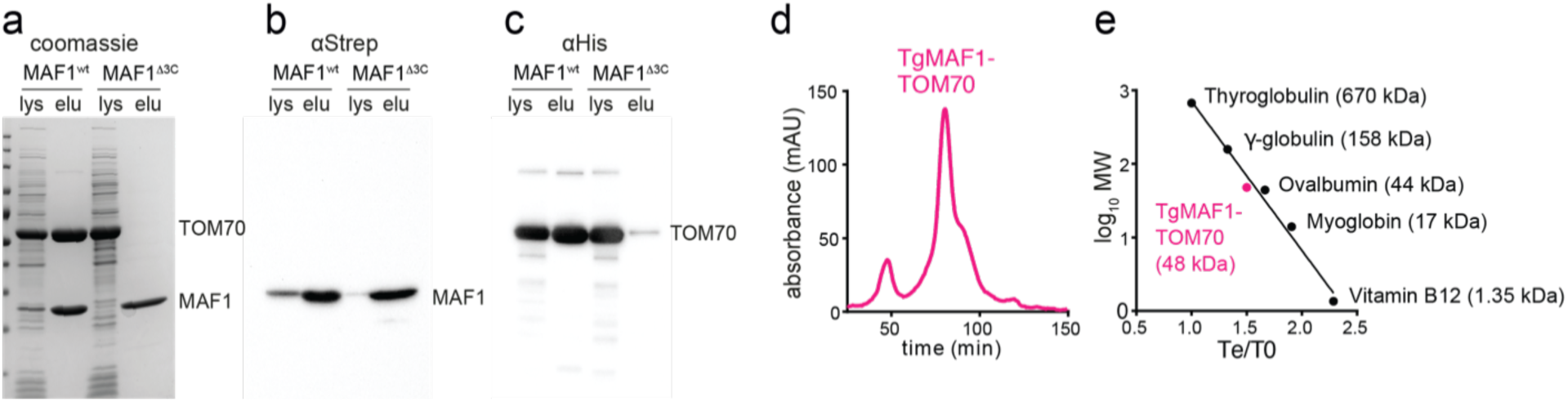
Isolation of HsTOM70-TgMAF1 complex. **a,** Coomassie of co-expression of His-TOM70^111-608^ and StrepII-TgMAF1^173-446^ or TgMAF1^Δ3C^ in *E. coli*. **b,** Immunoblot of (a) against StrepII-TgMAF1. **c,** Immunoblot of (a) against His-TOM70. **d,** Size-exclusion chromatography (SEC) of the purified TgMAF1-TOM70 complex. **e,** Molecular weight estimation of isolated TgMAF1-TOM70 by comparison with standard proteins using SEC.

**Extended Data Fig 2:**
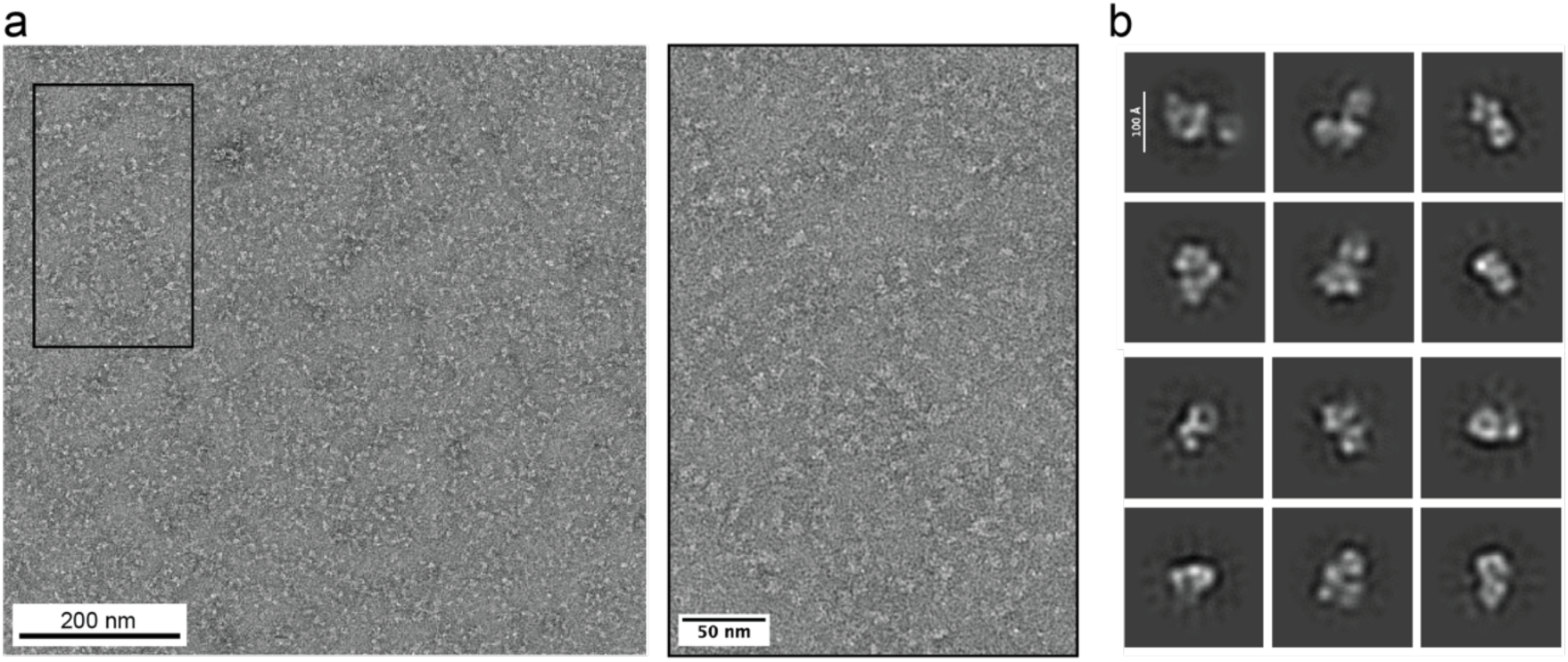
Negative staining analyses of the HsTom70-TgMAF1 complex. **a,** Representative negative stain micrograph of 200 nM HsTOM70-*Tg*MAF1 with zoomed inset; scalebars 200 nm and 50 nm **b,** 2D averages, showing mostly complexes of TOM70 with MAF1. Scalebar 10 nm.

**Extended Data Fig 3:**
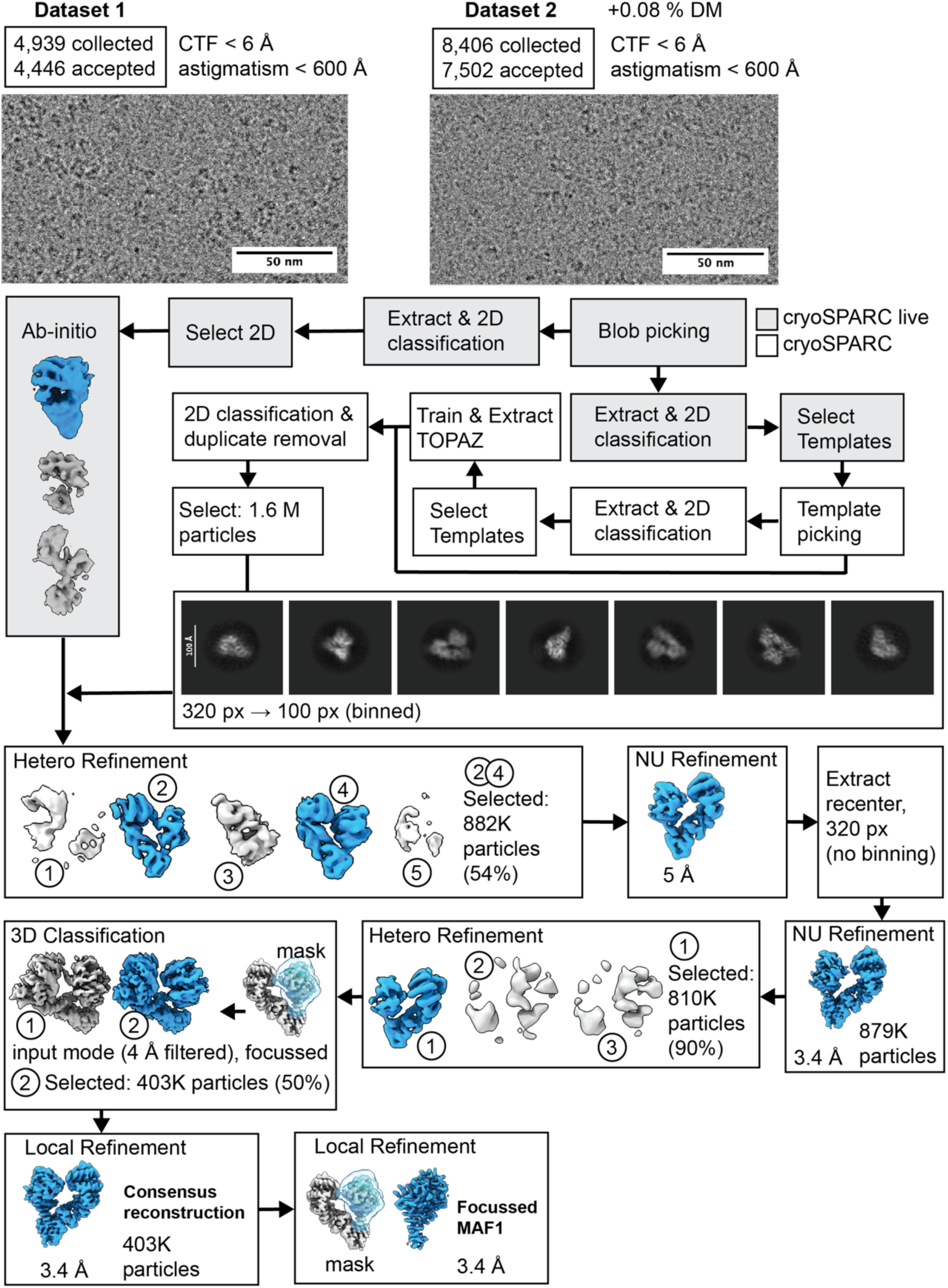
Cryo-EM data collection and image processing workflow. Representative micrographs and scale bar at 50 nm with their respective data collection statistics and applied selection criteria. Preprocessing, template creation for particle picking, and reconstruction of initial maps were generated in cryoSPARC live (shaded in gray). The combined and selected 2D class averages are shown with a 10 nm scale bar. All further processing steps were performed in cryoSPARC (no shading). Cryo-EM volumes in blue indicate selected volumes for further processing.

**Extended Data Fig 4:**
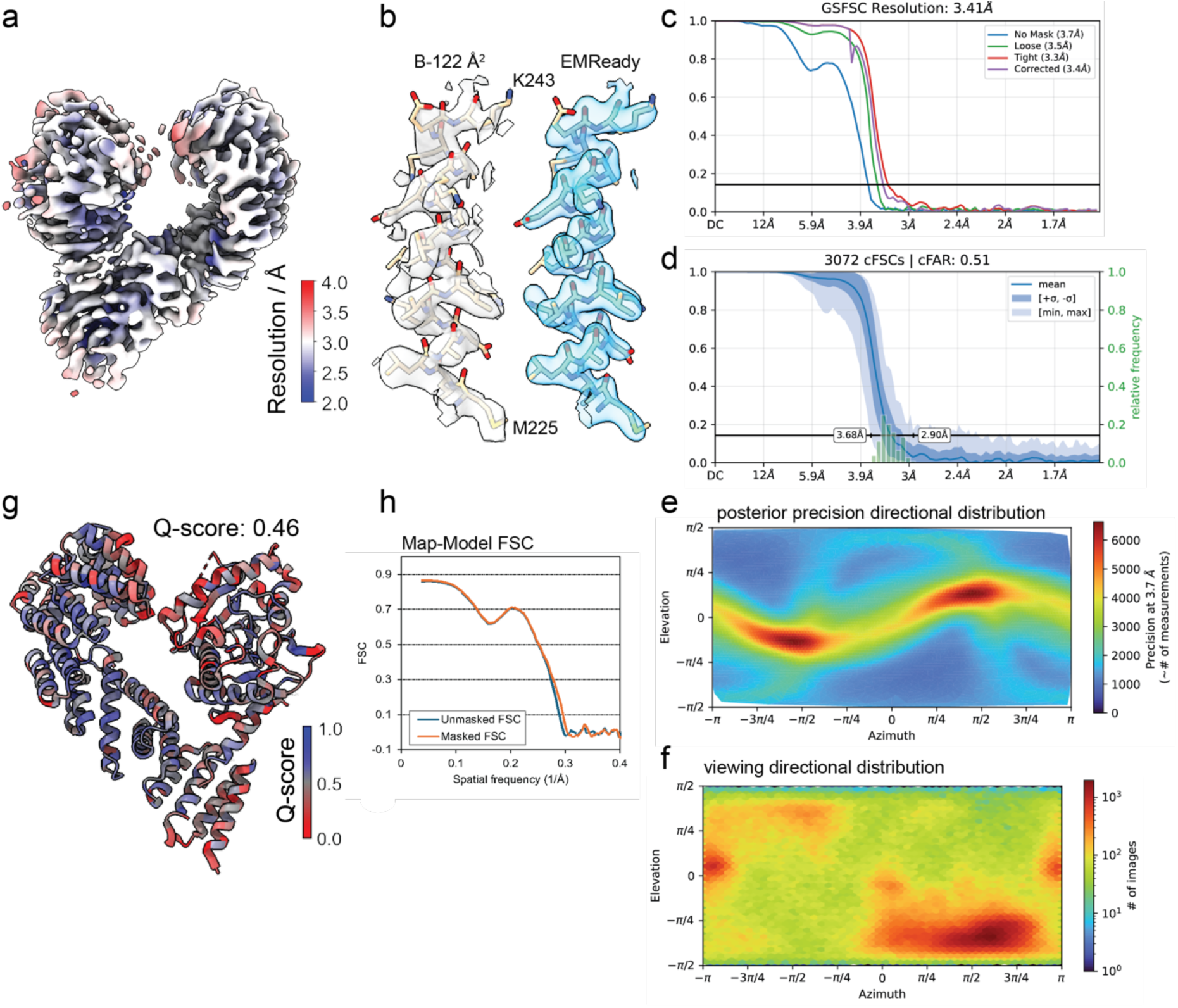
Validation of cryo-EM data. Comprehensive validation of the TOM70-TgMAF1 consensus reconstruction **a,** Local resolution distribution calculated with BlocRes in cryoSPARC. **b,** Map-model fit of the unmodified (grey) and EMReady-modified (blue) reconstruction. **c,** Gold-Standard Fourier-Shell Correlation reporting the overall resolution at FSC cutoff 0.134. **d,** Conical FSC report. cFAR values below 0.5 are considered anisotropic. **e-f,** Posterior precision of the directional distribution and viewing directions of the directional distribution. Precision assignments within one order of magnitude difference are considered balanced. **g,** Per-residue atom resolvability (Q-score) values were used to color the atomic model using ChimeraX. The median Q-score is 0.49, indicating moderate resolution and matches the reported resolution range. **h,** Map-to-Model FSC calculation from MolProbity within Phenix.

**Extended Data Fig 5:**
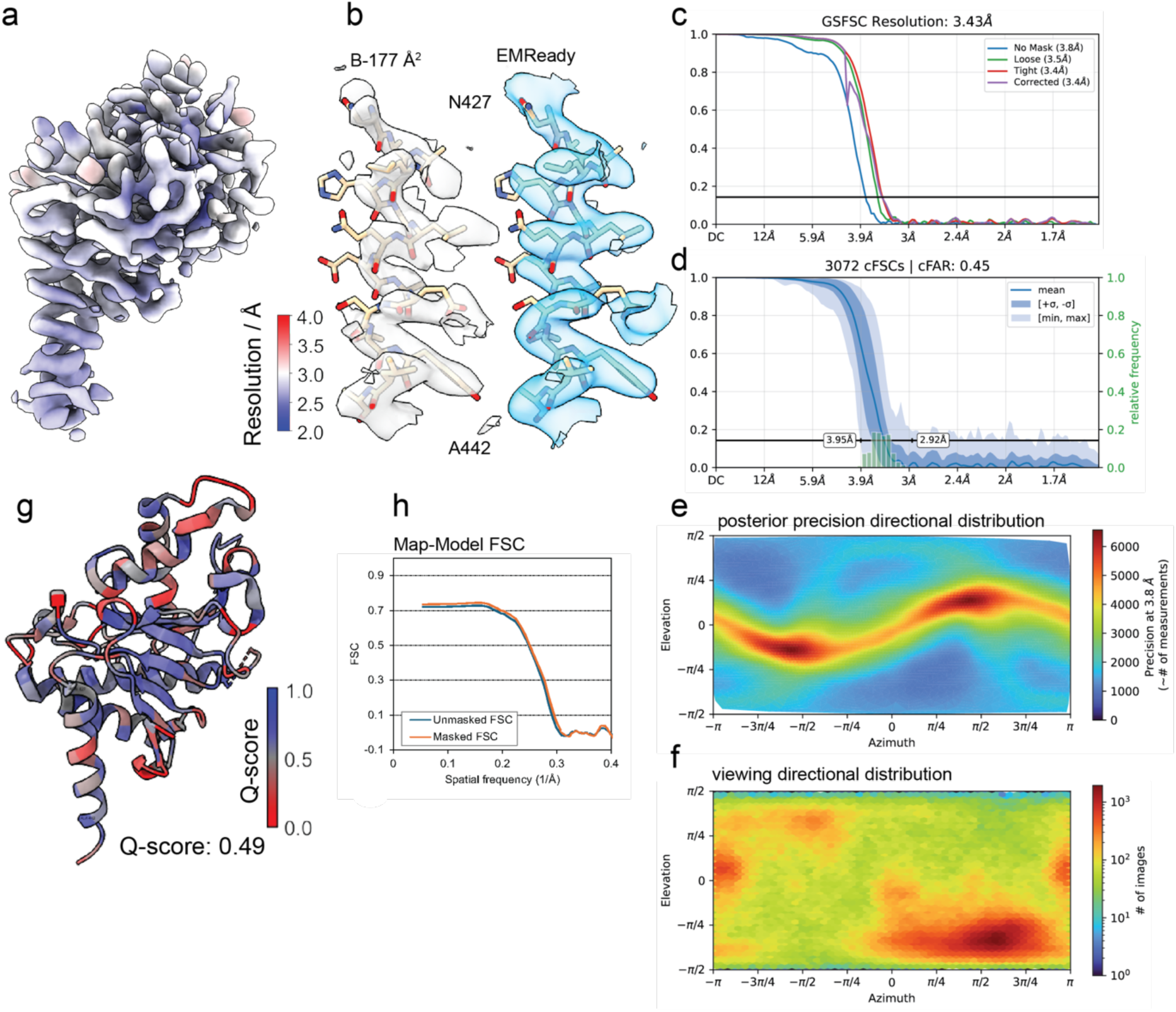
Validation of cryo-EM data. Comprehensive validation of the local MAF1 reconstruction **(a)-(h)** see Ext. Data 4 legend.

**Extended Data Fig 6:**
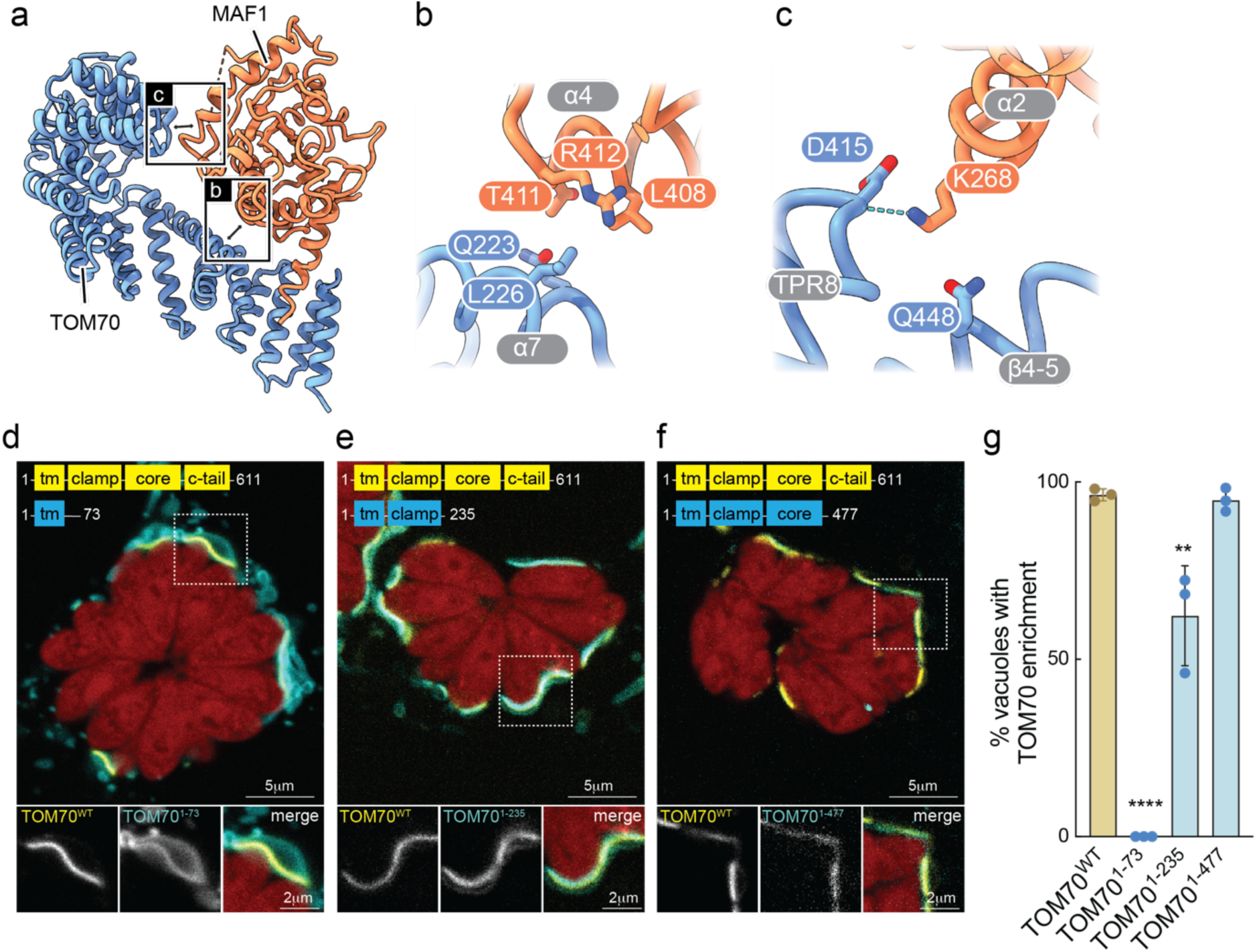
Head-to-Tail contact analysis of TOM70 TgMAF1. **a,** Model of TOM70 and MAF1 are colored in blue and orange, respectively. The inter-subunit contact is highlighted with a black box. **b,c** Involved secondary structures (grey) and contacting residues shown as sticks with polar interactions as blue dotted lines. **d-f,** Live cell images of cells stably expressing TOM70-GFP WT and the indicated TOM70 truncation mutant fused to BFP at 24 hours post infection with mCherry-expressing *Toxoplasma* (*Toxo-*mch). Scale bars 5μm and 2μm. **g,** Percentage of vacuoles positive for TOM70-sequestrations in images d-f; data are mean +/- SEM of more than 50 vacuoles counted from 3 biological replicates. ****p<0.001, **p<0.01 by one-way ANOVA, comparison to WT.

**Extended Data Fig 7:**
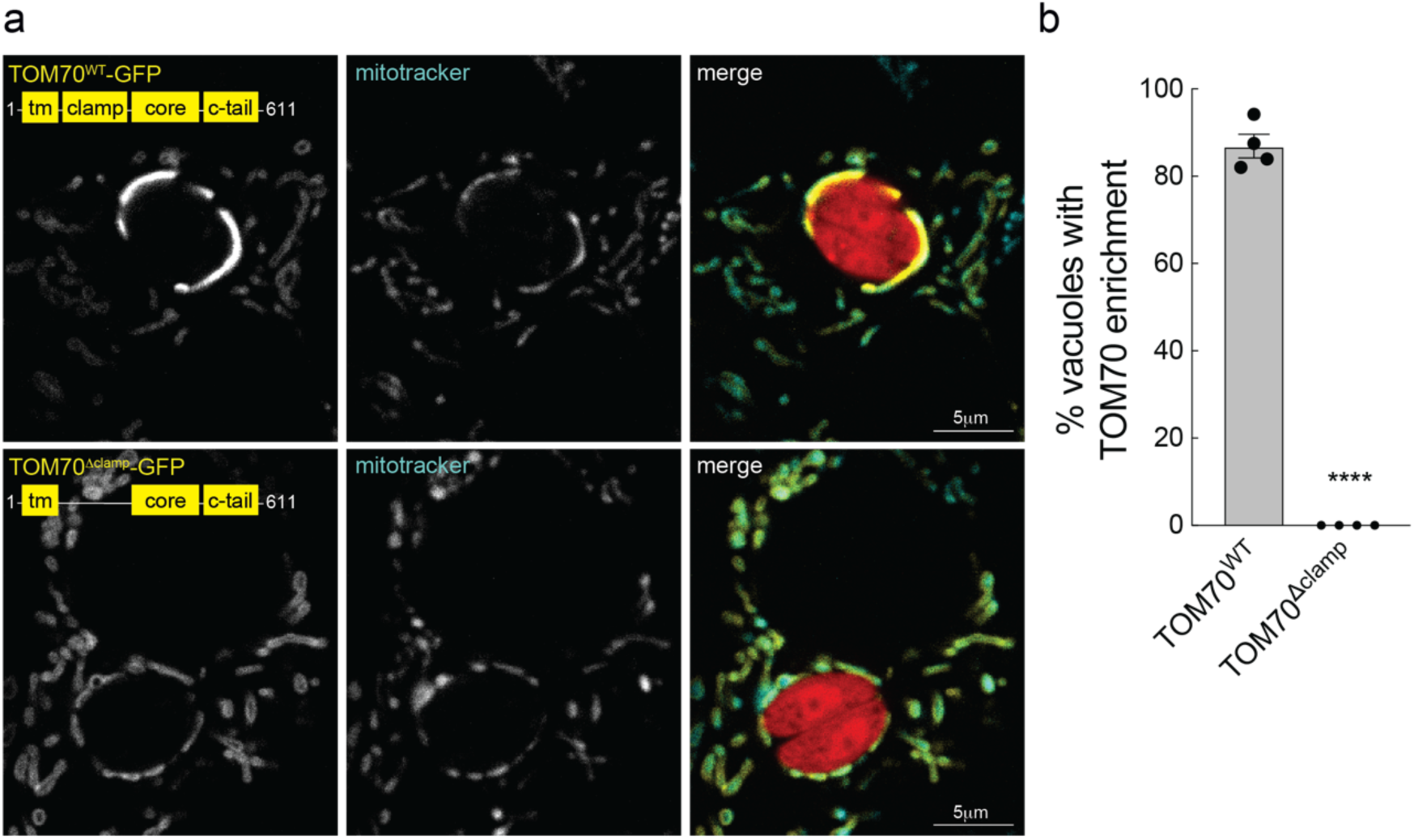
The clamp domain of TOM70 is required for its enrichment at host mitochondria-*Toxoplasma* PV contact sites. **a,** Live cell images of cells stably expressing TOM70^WT^-GFP or TOM70^Δclamp^-GFP and labeled with mitotracker at 24 hours post infection with mCherry-expressing *Toxoplasma.* Scale bars 5 μm. **b,** Percentage of vacuoles positive for TOM70-sequestrations in experiments as in A; data are mean ± SEM of more than 50 vacuoles counted from 3 biological replicates. ****p<0.001, **p<0.01 by one-way ANOVA, comparison to WT.

**Extended Data Fig 8:**
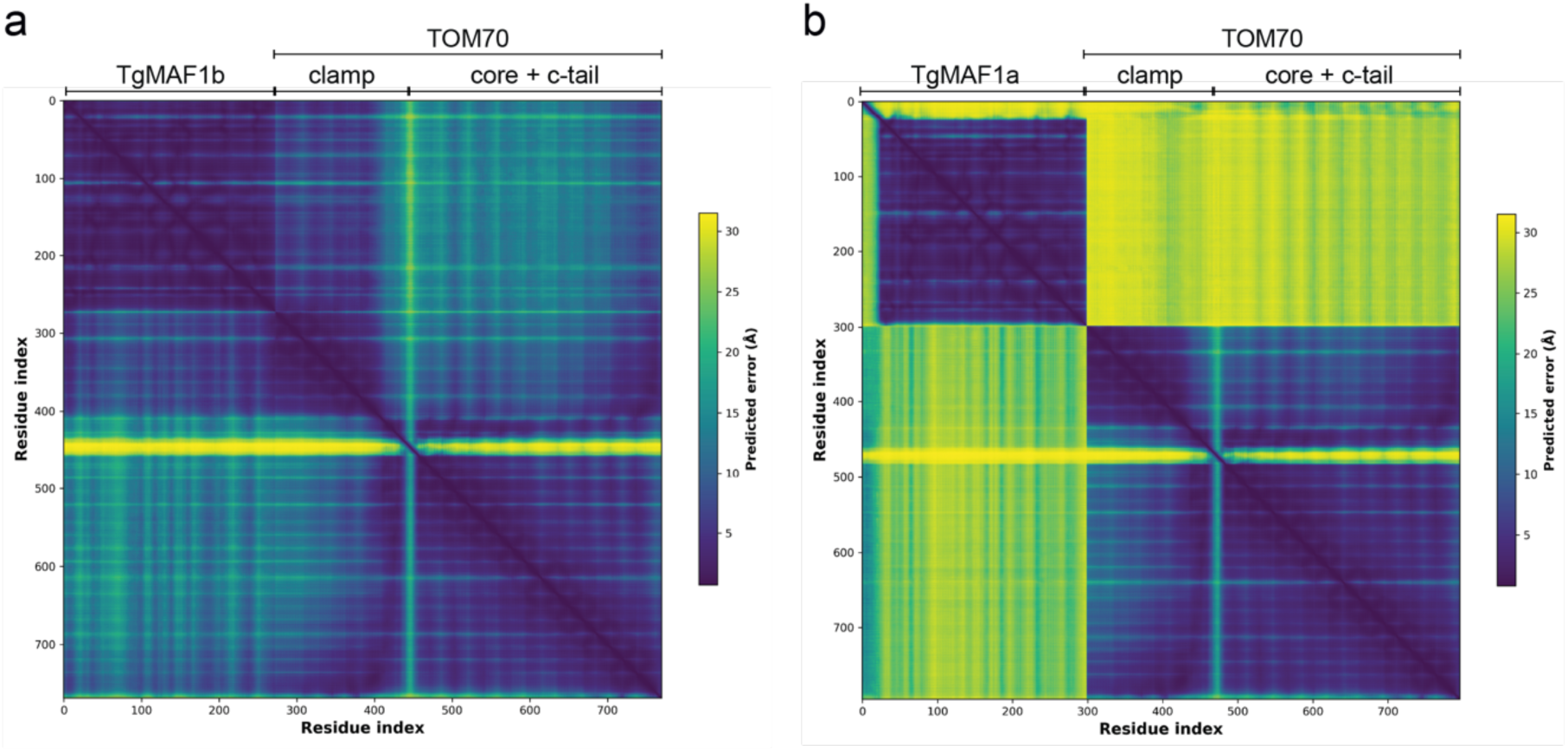
TgMAF1a is not predicted to interact with TOM70. AlphaFold2 predicted alignment error (PAE) plot of a putative complex between **a,** HsTOM70 and TgMAF1b and **b,** HsTOM70 and TgMAF1a.

**Extended Data Fig 9:**
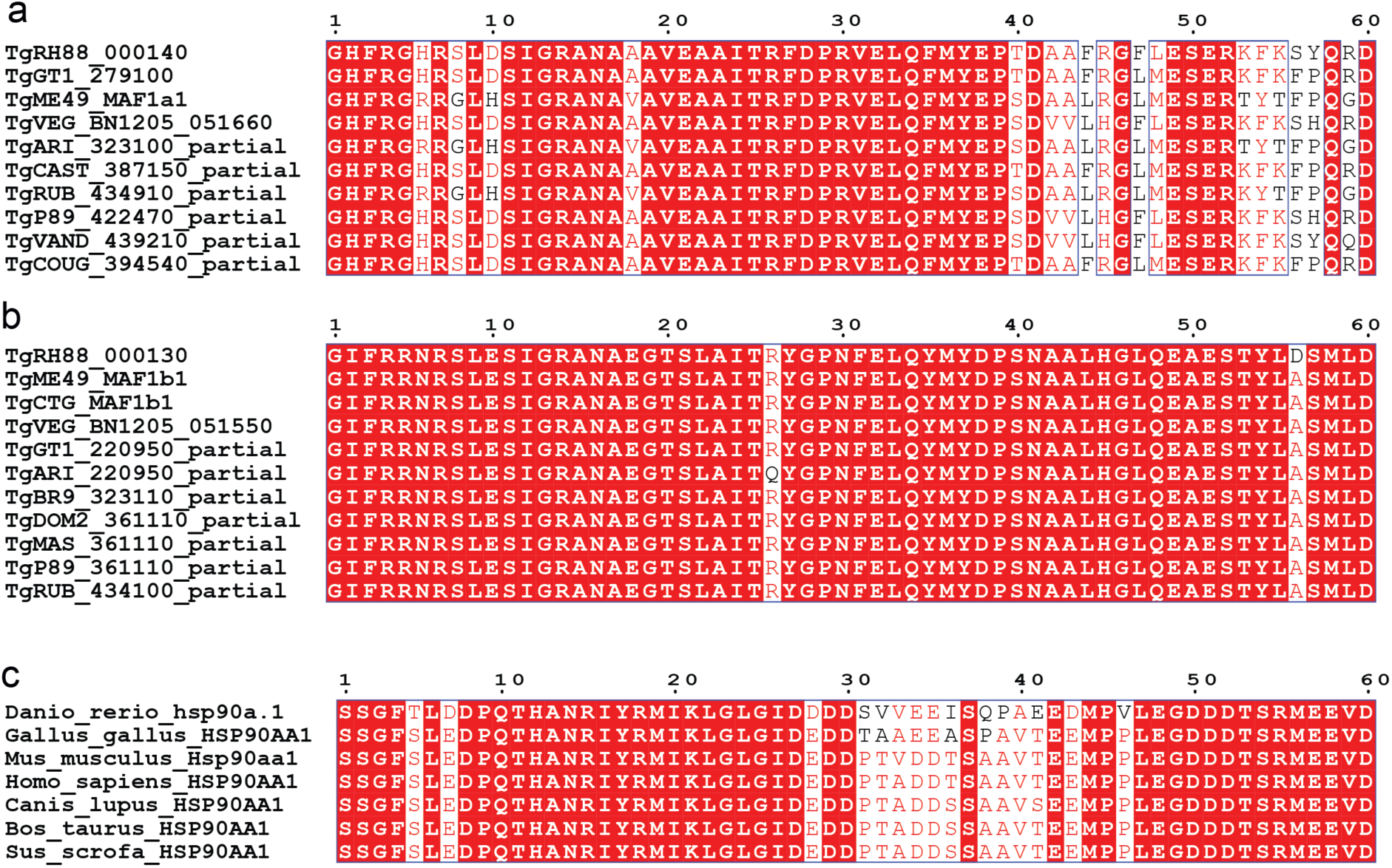
Alignment of TgMAF1a, TgMAF1b, and HSP90AA1 C-termini. **a,** Alignment of MAF1a and **b,** MAF1b termini from indicated *T. gondii* strains, and **c,** HSP90AA1 from the indicated species. Shaded red boxes indicates 100% identity; red text but not shaded indicates similarity in a group; blue boxes indicate similarity across groups. Alignments were visualized in ESPript 3.2.

**Extended Data Fig 10:**
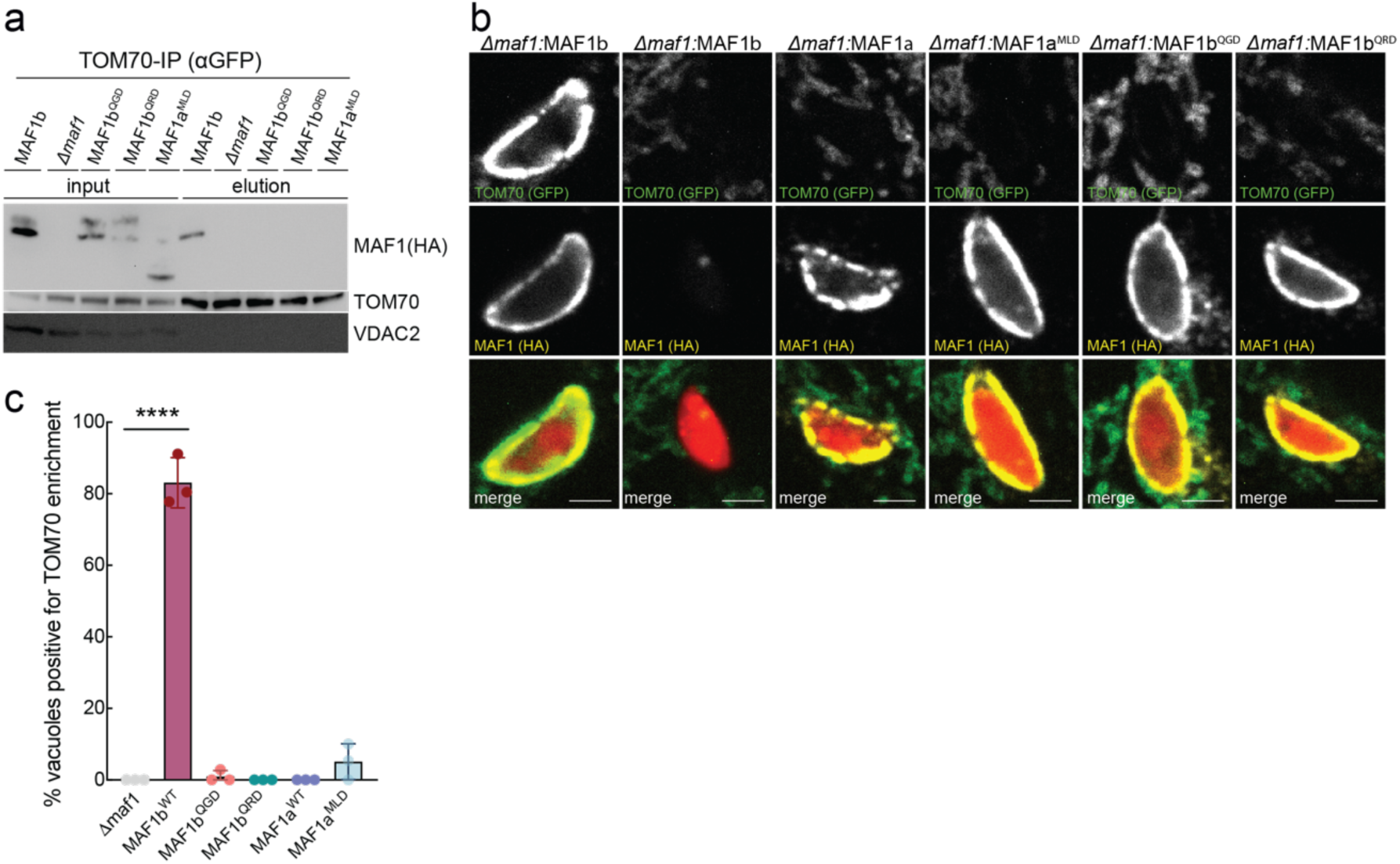
The TgMAF1 MLD motif is critical for its interaction with TOM70. **a,** anti-GFP IPs from murine embryonic fibroblasts (MEFs) stably expressing TOM70-GFP 24 hpi with indicated *Toxoplasma* strains and analyzed by means of immunoblotting (IB); TgMAF1 ∼60 kDa, TOM70 ∼72 kDa, VDAC2 ∼34 kDa. **b,** Immunofluorescence images of TOM70-GFP-expressing MEFs 5 hours post infection (hpi) with indicated mCherry-expressing *Δmaf1* mutants; scale bar 2 μm. (TgMAF1, HA; TOM70; GFP). **c,** Percentage of vacuoles positive for TOM70 enrichment in experiments as in (b); >30 vacuoles from n=3 biological replicates, ****p<0.001 by one-way ANOVA, comparison to *Δmaf1*.

**Extended Data Fig 11:**
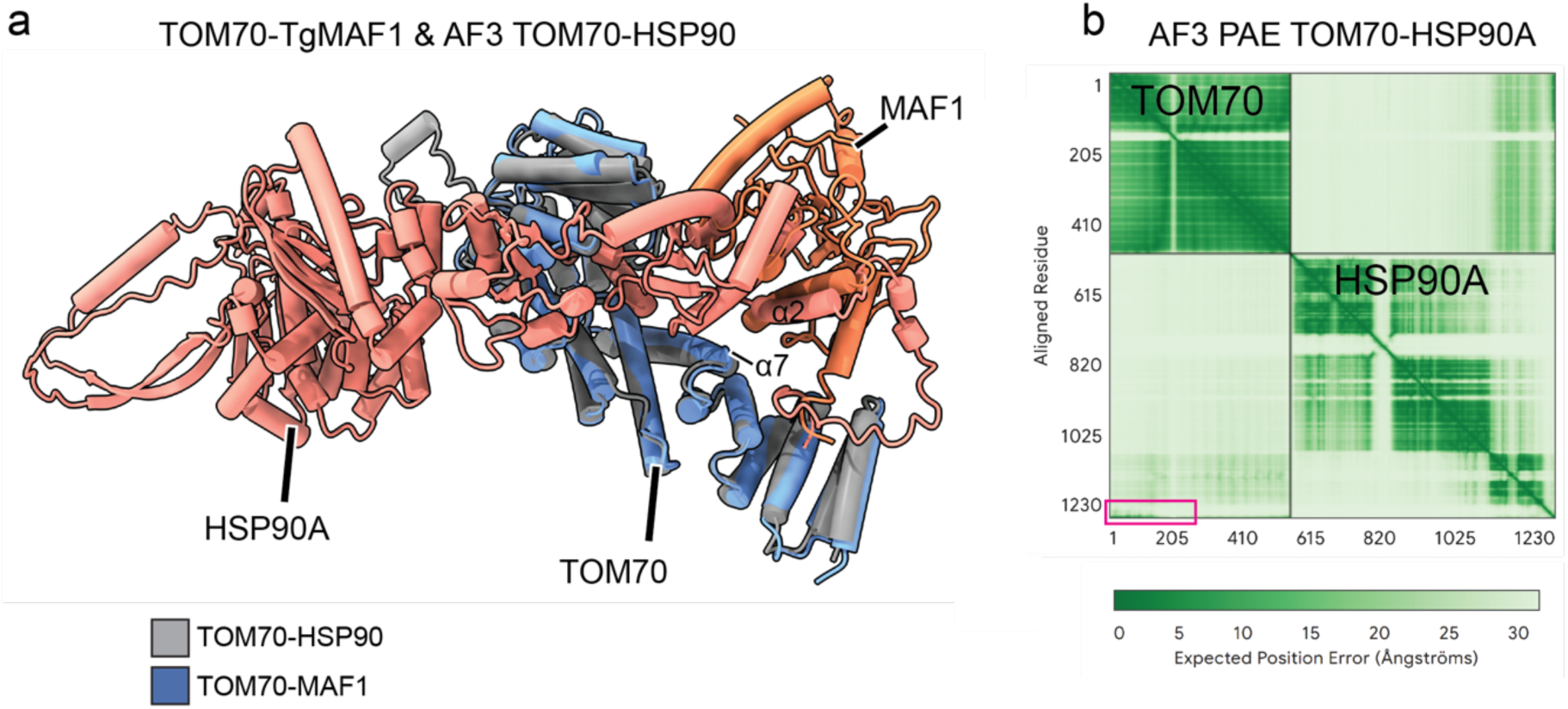
TgMAF1 and HSP90 are predicted to similarly engage TOM70. **a,** Superimposed AlphaFold3-predicted model of a putative complex consisting of TOM70 (Uniprot: O94826 111-611, gray) and HSP90A (Uniprot: P07900, salmon) with the experimental TOM70-MAF1 (blue and orange, respectively). **b,** Alphafold3 predicted aligned error matrix (PAE) of putative TOM70-HSP90A complex with chain borders annotated and C-terminus contact highlighted in a pink box.

**Extended Data Fig. 12:**
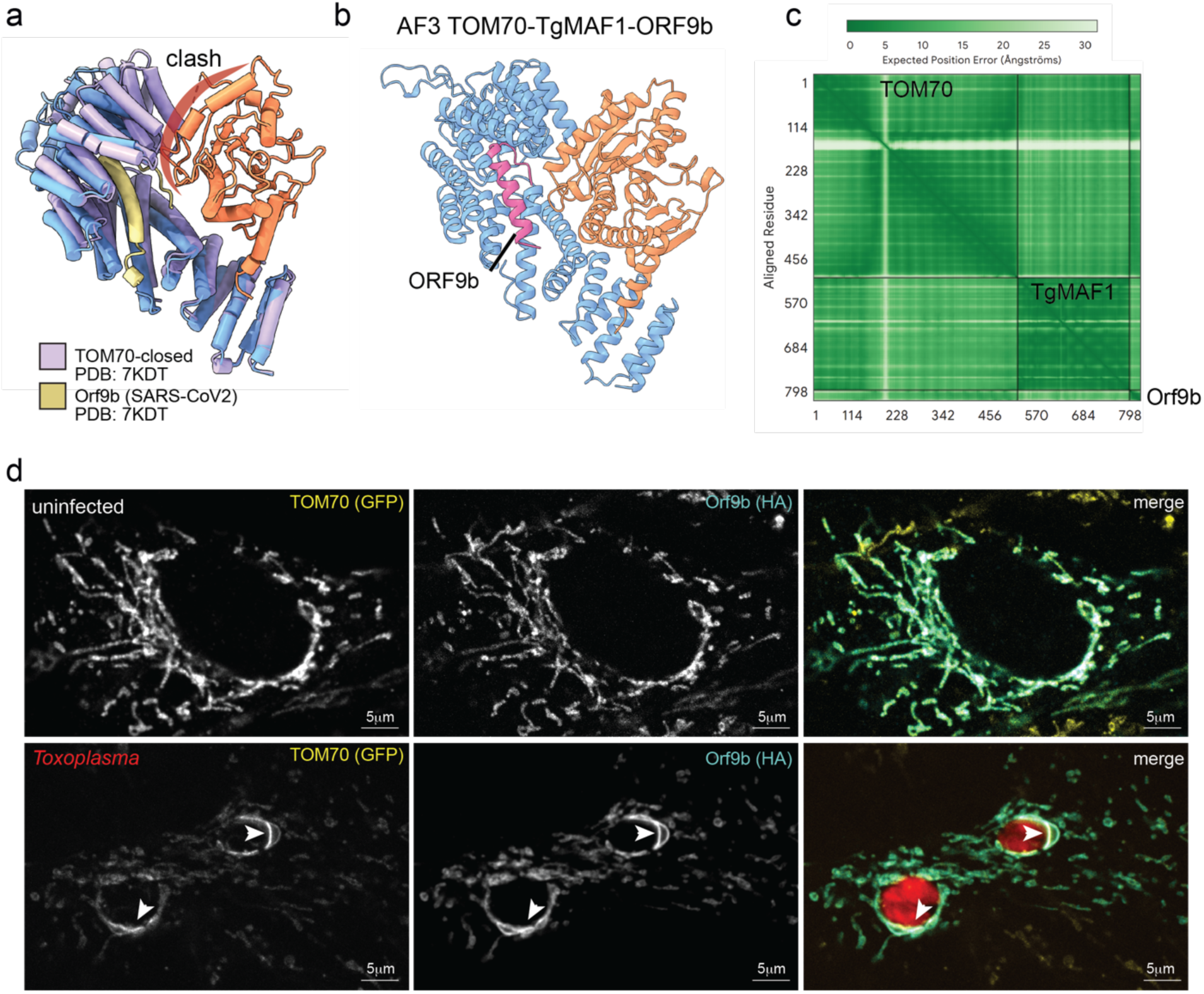
TgMAF1 binding to TOM70 does not impair Orf9b binding. **a,** Superimposition of the TOM70-SARS-CoV2-Orf9b complex (PDB: 7KDT, violet and yellow, respectively) and the TOM70-MAF1 complex, with clash of MAF1 and TOM70 c-tail indicated in red. **b,** AF3-predicted model of a putative complex consisting of TOM70 (Uniprot: O94826, residues 111-608, blue), Orf9b (Uniprot: P0DTD2, residues 44-70, pink), and TgMAF1 (Uniprot: A0A140H546, residues 173-446, orange). **c,** AF3-predicted aligned error matrix (PAE) of putative TOM70-Orf9b-TgMAF1 complex with chain borders annotated. **d,** Immunofluorescence images of MEFs expressing TOM70-GFP and Orf9b-HA mock-infected or infected with *Toxo-*mch and fixed at 5 hours post infection (hpi); scale bar 2 μm; Orf9b, HA; TOM70, GFP). Arrowheads indicate overlap of Orf9b and TOM70 sequestered at PVM-mitochondria interface.

**Extended Data Fig. 13:**
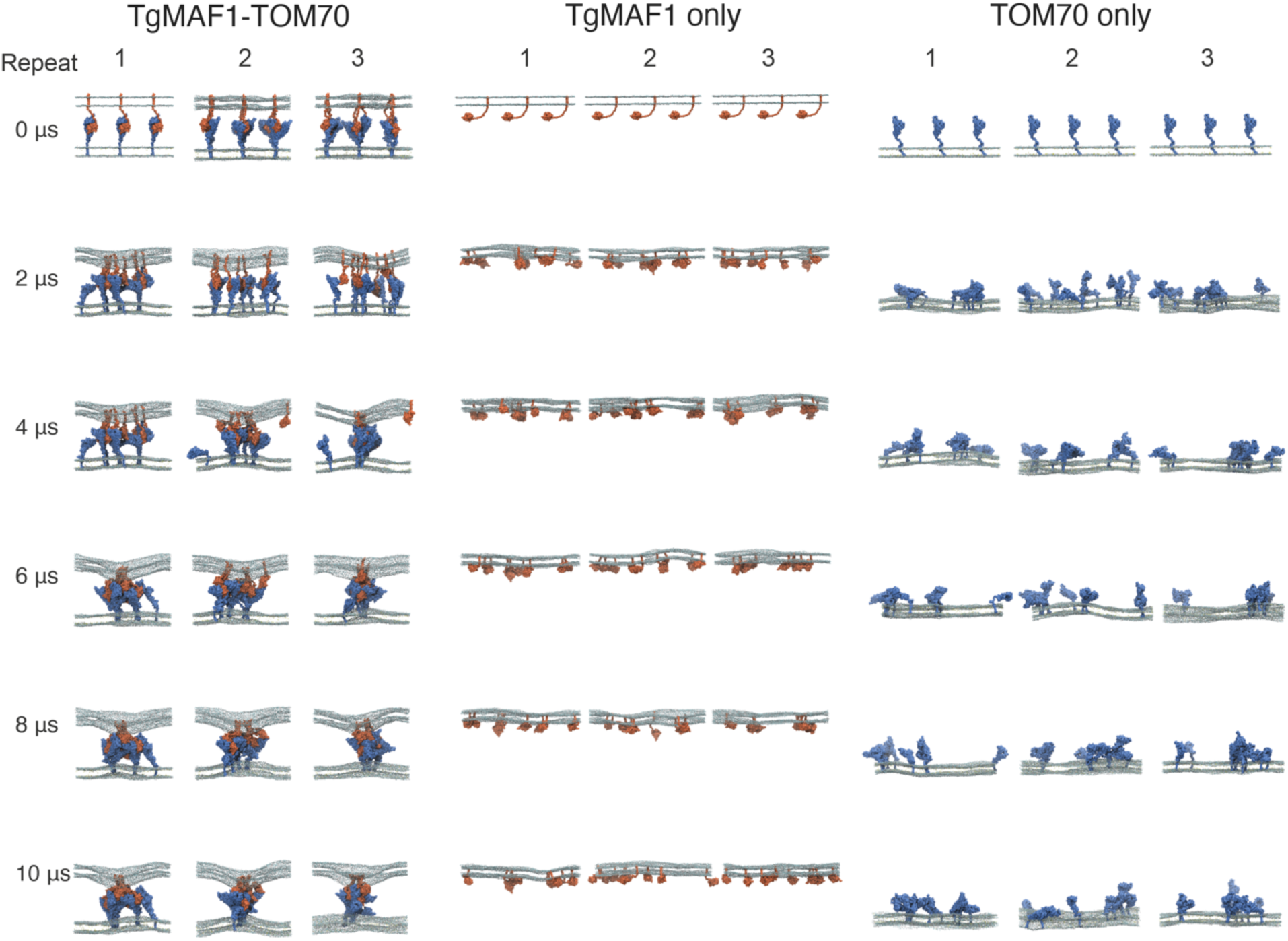
Overview of molecular dynamics simulations. Simulation snapshots of indicated proteins or complexes at 2 μs intervals with TgMAF1b (coral) and TOM70 (blue) shown in quicksurf representation and lipid headgroups shown as spheres: 1-palmitoyl-2-oleoylphosphatidylcholine (POPC, grey), 1-palmitoyl-2-oleoylphosphatidylethanolamine and POPE (light blue), and cholesterol (ROH, yellow).

**Extended Data Fig. 14:**
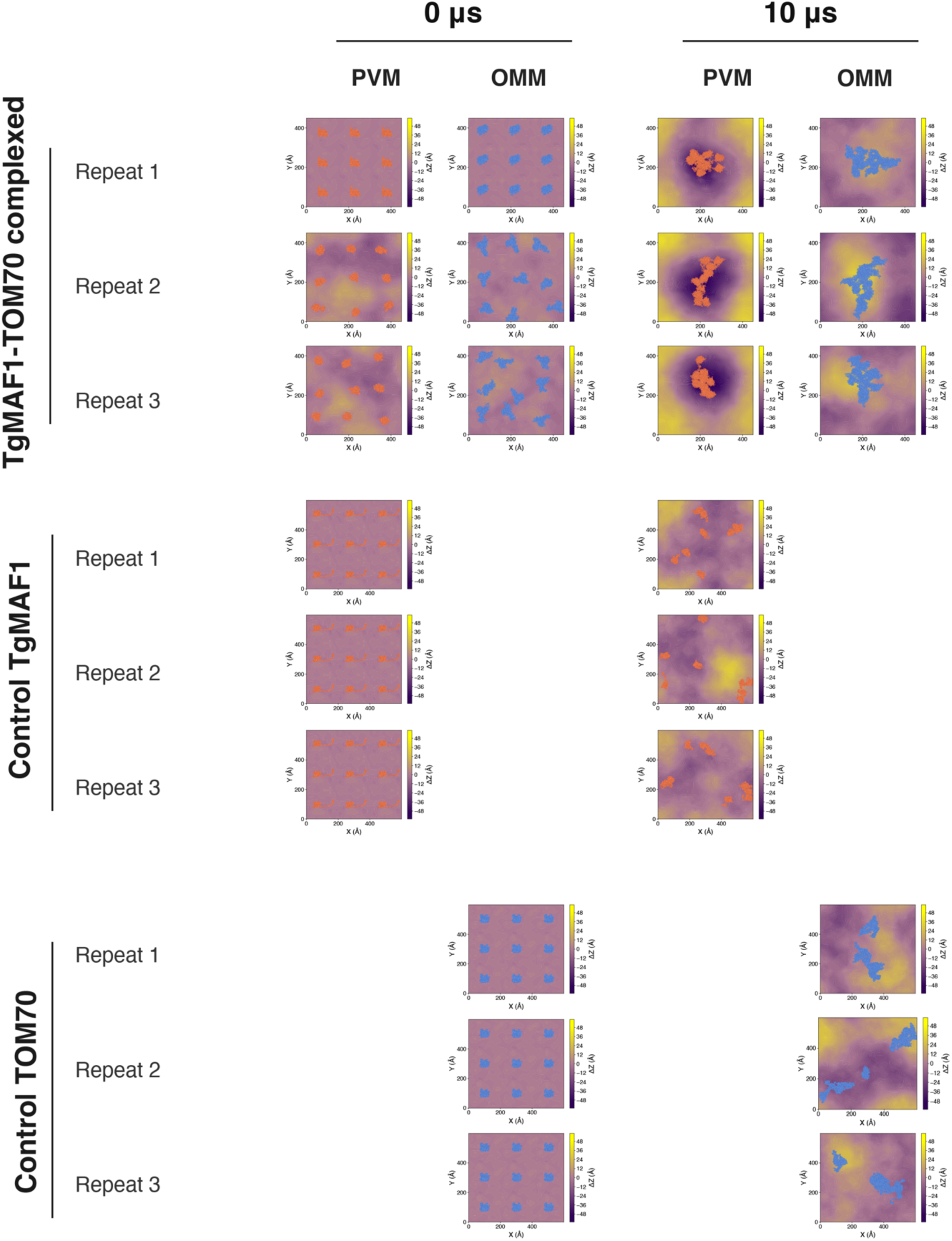
Instantaneous membrane curvature 2D heatmap plots of the PVM and OMM interfacial leaflets with instantaneous protein coordinates of indicated proteins or complexes (TgMAF1 (coral) and TOM70 (blue)) at 0 μs and 10 μs for all systems and repeats. The deviation from the mean Z height for the PO4 headgroups of the interfacial leaflet were plotted to represent the extent of membrane curvature in the simulation.

